# Integrated analysis of the aging brain transcriptome and proteome in tauopathy

**DOI:** 10.1101/2020.02.19.954578

**Authors:** Carl Grant Mangleburg, Timothy Wu, Hari K. Yalamanchili, Caiwei Guo, Yi-Chen Hsieh, Duc M. Duong, Eric B. Dammer, Philip L. De Jager, Nicholas T. Seyfried, Zhandong Liu, Joshua M. Shulman

**Affiliations:** Department of Molecular and Human Genetics, Baylor College of Medicine, Houston, TX 77030, USA; Medical Scientist Training Program, Baylor College of Medicine, Houston, TX, 77030, USA; Department of Pediatrics, Baylor College of Medicine, Houston, TX 77030, USA; Department of Neuroscience, Baylor College of Medicine, Houston, TX 77030, USA; Department of Biochemistry, Emory University School of Medicine, Atlanta, GA 30322, USA; Center for Translational & Computational Neuroimmunology, Department of Neurology and Taub Institute for the study of Alzheimer’s disease and the aging brain, Columbia University Medical Center, New York, NY 10032, USA; Cell Circuits Program, Broad Institute, Cambridge, MA 02142, USA; Department of Neurology, Emory University School of Medicine, Atlanta, GA 30322, USA; Jan and Dan Duncan Neurological Research Institute, Texas Children’s Hospital, Houston, TX 77030, USA; Department of Neurology, Baylor College of Medicine, Houston, TX 77030, USA

**Keywords:** MAPT, Tau, Alzheimer’s disease, transcriptome, proteome, inflammation, innate immunity

## Abstract

**Background:** Tau neurofibrillary tangle pathology characterizes Alzheimer’s disease and other neurodegenerative tauopathies. Brain gene expression profiles can reveal mechanisms; however, few studies have systematically examined both the transcriptome and proteome or differentiated Tau- versus age-dependent changes.

**Methods:** Paired, longitudinal RNA-sequencing and mass-spectrometry were performed in a *Drosophila* model of tauopathy, based on pan-neuronal expression of human wildtype Tau (Tau^WT^) or a mutation causing frontotemporal dementia (Tau^R406W^). Tau-induced, differentially expressed transcripts and proteins were examined cross-sectionally or using linear regression and adjusting for age. Hierarchical clustering was performed to highlight network perturbations, and we examined overlaps with human brain gene expression profiles in tauopathy.

**Results:** Tau^WT^ induced 1,514 and 213 differentially expressed transcripts and proteins, respectively. Tau^R406W^ had a substantially greater impact, causing changes in 5,494 transcripts and 697 proteins. There was a ~70% overlap between age- and Tau-induced changes and our analyses reveal pervasive bi-directional interactions. Strikingly, 42% of Tau-induced transcripts were discordant in the proteome, showing opposite direction of change. Tau-responsive gene expression networks strongly implicate innate immune activation, despite the absence of microglia in flies. Cross-species analyses pinpoint human brain gene perturbations specifically triggered by Tau pathology and/or aging, and further differentiate between disease amplifying and protective changes.

**Conclusions:** Our results comprise a powerful, cross-species functional genomics resource for tauopathy, revealing Tau-mediated disruption of gene expression, including dynamic, age-dependent interactions between the brain transcriptome and proteome.

## Background

The Microtubule Associated Protein Tau (MAPT/Tau) aggregates to form neurofibrillary tangle pathology in Alzheimer’s disease (AD) and other neurodegenerative tauopathies characterized by progressive cognitive and/or motor disability, including progressive supranuclear palsy (PSP), corticobasal degeneration, chronic traumatic encephalopathy, and certain forms of frontotemporal dementia (FTD) [1,2]. Rare mutations in the *MAPT* gene cause familial FTD, which is also characterized by prominent neurofibrillary tangle deposition [3–5]. Based on this genetic evidence, along with results from cellular and animal models [6, 7], Tau is a critical mediator of age-related neurodegeneration and a causal link among this diverse group of neurologic disorders. While the precise mechanisms of Tau-induced neuronal injury remain incompletely defined, progressive synaptic dysfunction and neuronal loss likely arises from a cascade of cellular derangements, including oxidative- and immune-mediated injury, altered proteostasis, and aberrant transcription and translation [6, 8].

RNA-sequencing (RNA-seq) makes possible comprehensive gene expression profiling of postmortem human brain tissue in AD and other tauopathies, providing a systems-level view of transcriptome perturbations accompanying neurodegeneration [9–13]. However, interpretation of differential gene expression analysis is hindered by a number of potential limitations. One major challenge arises from the recognition that the pathologic cascade in AD and related disorders initiates decades prior to onset of clinical manifestations [14, 15], whereas human brain expression profiles can only be generated cross-sectionally at the time of death. Indeed, it is essential to reconstruct the longitudinal, aging-dependent time-course of molecular derangements in order to pinpoint the earliest opportunities for intervention and to develop more effective biomarkers. Second, most brains from older persons with dementia show mixed pathologies at autopsy [16]. Therefore, it can be difficult to differentiate Tau-induced specific expression changes from those caused by other lesions (e.g. amyloid plaques, infarcts, etc.) or brain aging more generally. Third, among associated gene expression changes, it is important to identify those perturbations that are truly primary and therefore causal, rather than simply a consequence of disease. Lastly, emerging evidence suggests that transcription and translation are frequently discordant [17], making it important to consider both mRNA and protein changes to resolve many disease-associated expression signatures. While recent advances in mass-spectrometry permit deep surveys of protein expression, few studies have systematically profiled both the brain transcriptome and proteome in AD and related tauopathies [18,19].

By contrast with studies of human postmortem tissue, transgenic animal models of tauopathy readily permit controlled experimental manipulations to (i) define age-dependent changes, (ii) isolate the specific impact of Tau, and (iii) definitively establish causation. For example, RNA-seq in mouse transgenic models of tauopathy have highlighted early upregulation of inflammatory processes and downregulation of synaptic function genes preceding behavioral phenotypes, and suggest Tau-specific impact on microglial and neuronal function [20–23]. Expression of human *MAPT* in the nervous system of the fruit fly, *Drosophila melanogaster*, recapitulates many key features of tauopathies, including misfolded/hyperphosphorylated Tau, age-dependent synaptic dysfunction and neuronal loss, and reduced survival [6, 24]. Importantly, *Drosophila* permits high-throughput genetic manipulation, and these models have been successfully deployed for enhancer-suppressor screens [25–27]. The results highlight many promising modifiers of Tau-mediated neurodegeneration, including genes that overlap with human AD susceptibility loci [28–30]. Prior gene expression studies in fly tauopathy models have been limited by incomplete coverage [31] or cross-sectional design [32], and none have coupled analyses of both transcripts and proteins. We have analyzed longitudinal, paired transcriptome and proteomes from control flies and following pan-neuronal expression of either wildtype or mutant forms of human Tau. We identify Tau-induced patterns of differential expression that are robust to adjustment for aging, and we integrate our results with complementary expression profiles from human brains affected by tauopathy and known genetic modifiers of Tau neurotoxicity.

## Results

### Paired Tau transcriptomes and proteomes in *Drosophila*

Longitudinal, parallel RNA-seq and mass-spectrometry proteomics were performed in controls (*elav-GAL4*) and in flies with pan-neuronal expression of human wildtype (*elav>Tau*^*WT*^) or mutant Tau (*elav>Tau^R406W^*). The transgenic genotypes and age timepoints (1-, 10-, and 20-days) selected for this analysis have been extensively characterized in prior published work [6, 24], and we confirmed that Tau^WT^ and Tau^R406W^ are expressed at similar levels (Additional file 1: Figure S1). Overall, 17,104 transcripts and 2,723 proteins were detected. We first analyzed our results cross-sectionally, highlighting those transcripts or proteins significantly differentially expressed (FDR < 0.05) at each timepoint (Table 1; Additional File 2: Table S1). Overall, Tau^WT^ altered expression of 1,514 transcripts and 213 proteins. At each age examined, Tau^R406W^ induced a ~4- to 7-fold greater number of differentially expressed genes than Tau^WT^, highlighting 5,494 transcripts and 697 proteins. There was substantial overlap between the Tau^WT^ and Tau^R406W^ gene expression profiles, with 70% of Tau^WT^-associated transcripts showing consistent changes in Tau^R406W^ flies (65% of proteins) (Additional File 1: Figure S2). Overall, nearly equal numbers of up- or down-regulated, differentially expressed genes were detected in the Tau transcriptome; whereas in the proteome, Tau-induced gene up-regulation was more common by a factor of 2, which may reflect reduced assay sensitivity for proteins with low expression levels.

**Table 1:** Tau-triggered differentially expressed genes. Differentially-expressed transcripts (and proteins, in parentheses) are indicated based on cross-sectional comparisons in 1-, 10-, or 20-day-old *elav>Tau*^*WT*^ or *elav>Tau*^*R406W*^ animals and controls. Based on PCA analysis [40], the decrease in differentially expressed transcripts at day 10 in *Tau*^*R406W*^ flies is likely due to sample heterogeneity. The total number of unique differentially expressed transcripts/proteins from the cross-sectional analyses are also indicated, along with complementary results from the joint regression model including all longitudinal data and adjusting for age. Statistical analysis was based on a Wald test (FDR<0.05). See Additional File 2: Tables S1 and S4 for complete results.

|  | Day 1 | Day 10 | Day 20 | Total |  |
| --- | --- | --- | --- | --- | --- |
|  |  |  |  | cross-sectional | age-adjusted* |
| **Tau<sup>WT</sup>** | 491 (54) | 431 (143) | 1,096 (76) | 1,514 (213) | 1,653 (123) |
| **Tau<sup>R406W</sup>** | 3,179 (97) | 1,616 (173) | 4,087 (581) | 5,494 (697) | 4,992 (503) |

As in human tauopathy, the neurodegenerative phenotypes manifested by Tau transgenic flies are progressive with aging [24]. Consistent with this, we observed age-dependent differences in the number and identity of differentially expressed genes across the timepoints examined (Additional File 1: Figure S2). For example, only a minority (~9%) of transcripts from Tau^WT^ flies were consistently, differentially expressed at all 3 timepoints. The profound impact of aging on the *Drosophila* brain transcriptome and proteome is readily apparent in transcriptome-wide heatmaps (Fig. 1a,b) and unsupervised clustering analysis further highlights age as a major driver of gene expression differences among samples (Additional File 1: Figure S3). Indeed, among control animals, we documented age-related, differential expression of 6,742 transcripts and 1,155 proteins (Table 2 and Additional File 2: Tables S2, S3), and similar changes were seen in analyses of aged *Tau* animals (within genotype comparisons of data from different timepoints). Strikingly, approximately 70% of Tau-triggered transcripts overlap aging-associated gene expression changes. These data highlight an intimate connection between aging and Tau-mediated perturbations in gene expression.

**Figure 1:**
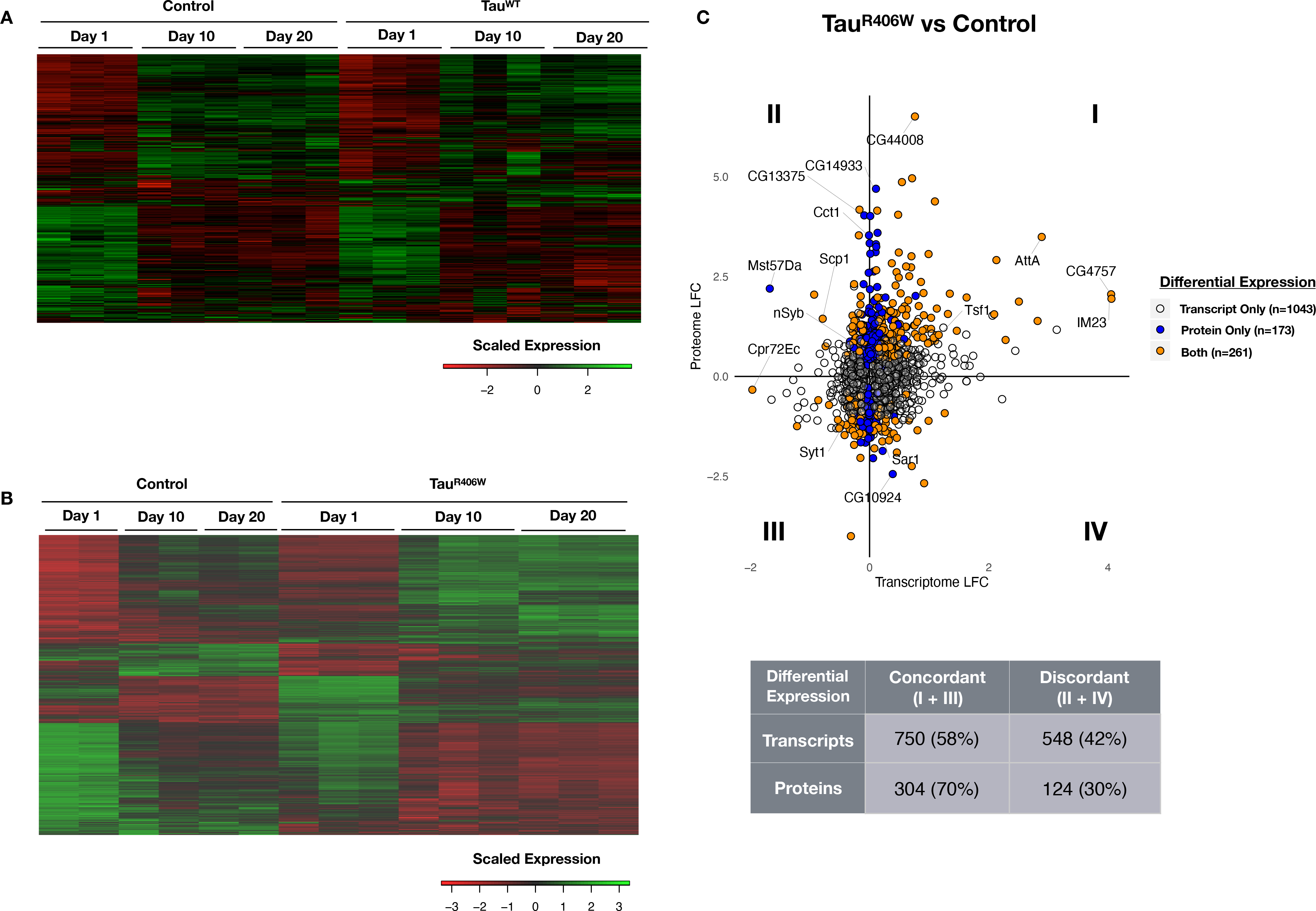
Tau-triggered differentially expressed genes. **a** Gene expression heatmap showing replicate samples from control flies (*elav*, n=3) and *elav > Tau*^*WT*^ (Tau^WT^, n=3) grouped by age (1-, 10-, and 20- days). Columns denote individual samples. Rows consist of clustered, normalized expression values for all differentially-expressed transcripts (n=1653, FDR<0.05) based on the joint regression model adjusting for age. Each column represents an individual sample. In both control and Tau^WT^ animals, age is the dominant driver of gene expression patterns. **b** Gene expression heatmap showing replicate samples from batch-matched control flies (*elav*, n=2) and *elav > Tau*^*R406W*^ (Tau^R406W^, n=3). Rows consist of clustered, normalized expression values for all differentially-expressed transcripts (n=4992, FDR < 0.05) based on the joint regression model adjusting for age. While age remains a major driver, Tau^R406W^ has a more substantial and appreciable impact on expression pattern compared with Tau^WT^ (a, above). **c** Plot (top) showing Tau^R406W^-triggered log_2_ fold-change (LFC) in the transcriptome and proteome. The plot only includes those genes detected as both transcripts and proteins and also differentially expressed (n=1477, FDR < 0.05), based on the joint regression model including longitudinal data and adjusting for age. Colors denote whether the gene was differentially expressed in the transcriptome (unfilled), proteome (blue), or both (orange). Quadrants I and III include gene expression changes that are concordant (same direction) at the transcript and protein level; whereas quadrants II and IV depict discordant changes. A substantial proportion of differentially-expressed transcripts or proteins are discordant (table, bottom). See Fig. 2 and Additional File 1: Figure S5 for selected examples (labeled).

**Table 2:** Aging-triggered differentially expressed genes. In *Tau*^*R406W*^ animals, aging is associated with an increased number of differentially-expressed transcripts but a decreased number of proteins. Differentially-expressed transcripts and proteins were determined by comparing aged animals, stratified by genotype, analyzing *elav* (control) or *elav>Tau*^*R406W*^ animals separately. The total number of unique, differentially-expressed transcripts or proteins are shown based on the union of 3 comparisons (1- vs. 10- days, 10- vs. 20-days, and 1- vs. 20-days). Statistical analysis was based on a Wald test (FDR<0.05). See Additional File 2: Tables S2 and S3 for complete results.

|  | Control | Tau<sup>R406W</sup> | Change |
| --- | --- | --- | --- |
| **Transcripts** | 6742 | 7970 | +18% |
| **Proteins** | 1155 | 258 | -78% |

**Figure 2:**
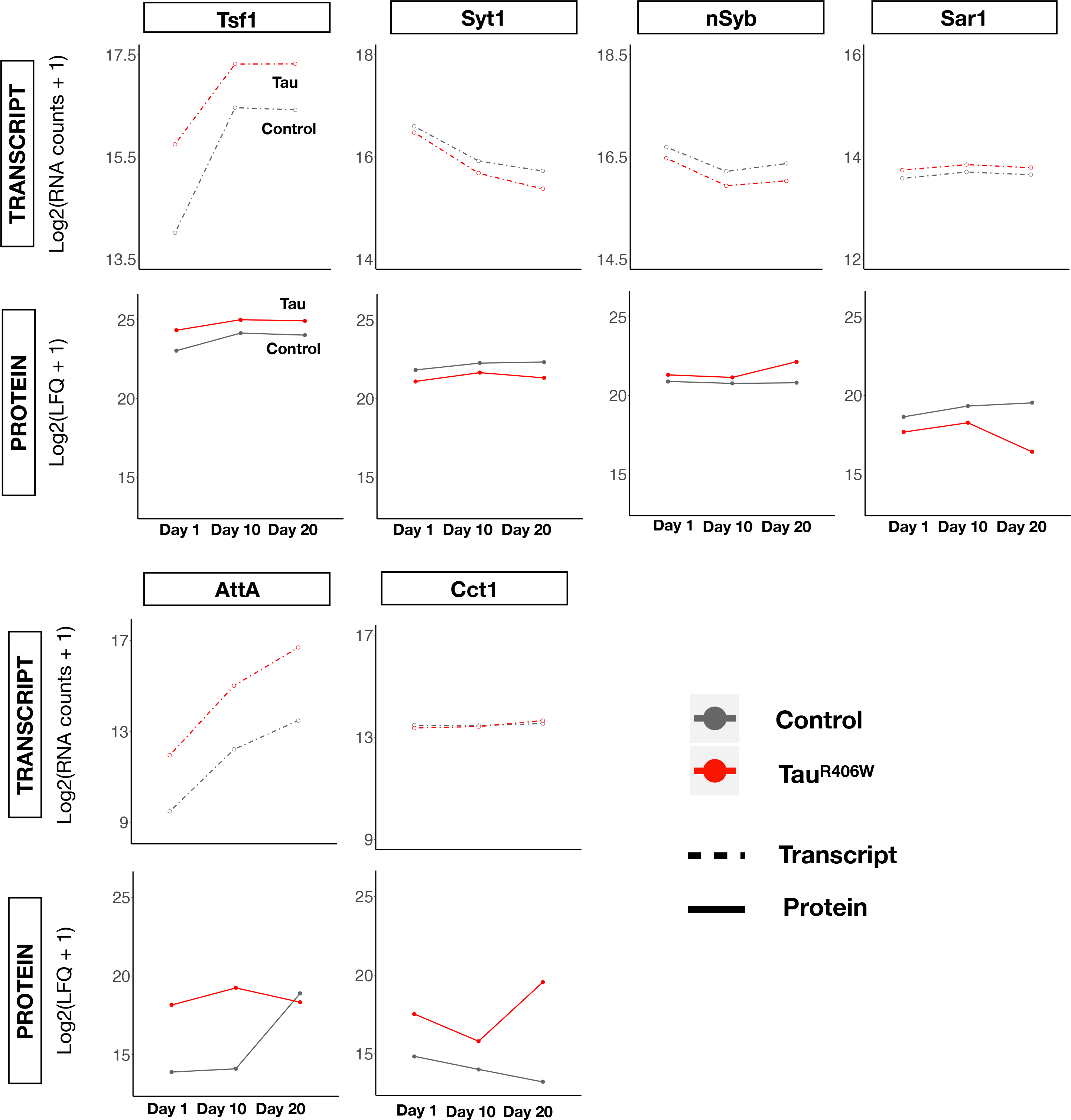
Examples of Tau-induced changes in the transcriptome and proteome. log_2_-transformed expression of selected genes in *elav>Tau*^*R406W*^ (Tau, red) and *elav* (Control, gray) is shown for transcriptome (depth normalized counts) and proteomes (normalized label-free quantification intensity (LFQ)). Genes were selected to be representative within our dataset and are all differentially expressed (FDR < 0.05) in both the transcriptome and proteome, based on the joint regression model including all longitudinal data and adjusting for age. CCT1 is only differentially-expressed at the protein level. *AttA* and *CCT1* transcripts (bottom) are plotted on a different scale than the other examples due to the increased dynamic range of changes (for *AttA*). Additional example plots can be found in Additional File 1: Figure S5.

### Integrated longitudinal analysis of differentially expressed transcripts and proteins

In order to identify the most robust, Tau-induced expression changes independent of aging, we used linear regression and considered all longitudinal data in a joint model, including a covariate to adjust for age. In separate analyses of Tau^WT^ and Tau^R406W^, we identify 1,653 and 4,992 significant differentially expressed transcripts, respectively (Table 1 and Additional File 2: Table S4). The same approach was applied to the longitudinal proteomic data. To better understand the joint impact of Tau on the transcriptome and proteome, we next examined those genes detected by both the RNA-seq and mass-spectrometry assays (n=2,395 and 2,423 for Tau^WT^ and Tau^R406W^, respectively). For this and subsequent analyses, we focus on the Tau^R406W^ dataset given the more substantial number of differential expression changes (analyses of Tau^WT^ are included as supplemental data and show consistent results). Remarkably, among 1,309 Tau^R406W^-triggered, differentially expressed transcripts with corresponding proteome measurement, only 58% show concordant changes in the proteome (same direction of change, regardless of significance) (Fig. 1c and Additional File 1: Figure S4). These data indicate that for a substantial proportion of transcriptional changes (42%), the behavior of corresponding proteins is discordant. Consistent with this, out of 503 significant, differentially expressed proteins induced by Tau^R406W^, 272 (48%) are unique to the proteome (e.g. either non-significantly changed or not detected in the transcriptome).

Tau-mediated perturbations of the transcriptome and proteome are readily appreciated in an integrated plot (Fig. 1c) including all significant, differentially expressed transcripts and/or proteins, and representative examples discussed below are highlighted in Fig. 2 (see also Additional File 1: Figure S5). Concordant activation or suppression of gene activation, respectively, is represented in the upper right (I) and lower left quadrants (III) of the plot. Many Tau-responsive genes show highly consistent and concordant expression changes in transcripts and proteins. *Transferrin 1* (*Tsf1*) encoding an iron-binding protein induced during the *Drosophila* innate immune response [33] is strongly activated by Tau^R406W^, showing similar ~2-fold increase in both the transcriptome and proteome, and these changes are largely consistent in 1-, 10-, and 20-day-old animals. Reciprocally, *Synaptotagmin-1* (*Syt1*), encoding the essential calcium sensor for synaptic vesicle release and neurotransmission [34], is decreased 10% at the transcript level and 40% at the protein level, and this result agrees with prior targeted studies of synaptic proteins in Tau^R406W^ flies [35]. By contrast, Tau-triggered gene expression changes that are discordant between the transcriptome and proteome occupy the upper left (II) and lower right (IV) quadrants of the plot (Fig. 1c). For example, Synaptobrevin (nSyb), which participates in synaptic vesicle fusion and release [36], is increased in Tau flies, whereas *nSyb* transcripts are reciprocally decreased. Alternatively, in the case of *Sar1*, encoding a GTPase involved in endocytic trafficking [37], we detect a Tau-associated increase in transcripts, whereas Sar1 protein is decreased. Such discordant changes may suggest the possibility of feedback regulation between the transcriptome and proteome. In other cases, we detect significant Tau-induced changes in the proteome without a corresponding change in transcript levels. One such example is *CCT1*, encoding a cytosolic chaperone implicated in cytoskeletal regulation and nerve injury response [38].

As suggested above, aging has a profound impact on brain gene expression and frequently modifies the impact of Tau, sometimes with divergent consequences in the transcriptome and proteome. For example, aging is associated with a substantial increase in *Tsf1* transcript expression in both Tau and control animals (~3- and 6-fold, respectively), whereas protein expression appears stable over the same timecourse. The immune response gene, *Attacin-A* (*AttA*), encoding an antimicrobial peptide, provides another striking example. RNA-seq reveals a consistent aging- and Tau-associated increase in *AttA* transcripts. However, the substantial Tau-associated increase observed in the proteome of 1-day-old flies is attenuated during aging and no longer detected by 20-days (genotype × age interaction, p=3.78×10^−3^). The sharp increase of AttA in wildtype flies with aging was previously reported and linked to neuronal maintenance [39]. Notably, in our age-adjusted joint model, only 35% of Tau-triggered differentially expressed transcripts were fully independent of aging. By contrast, the majority (65%) were both Tau- and aging-associated gene expression changes (Additional File 1: Figure S6). Given the pervasive impact of aging, we again carefully considered all aging-associated changes, focusing on relative changes across the transcriptome and proteome, as well as a potential interaction with Tau-mediated toxicity (Table 2 and Additional File 2: Table S2, S3). Interestingly, in Tau^R406W^ flies we note an ~18% increase in aging-associated transcripts, with 7,970 genes affected (versus 6,742 in controls). However, within the proteome, the reverse pattern is seen with only 258 age-associated protein changes detected (versus 1,155 in controls), representing a 78% reduction, and potentially consistent with reports of Tau-induced translational dysregulation [40–42]. A similar trend for the proteome is observed in Tau^WT^ animals; although, the magnitude of changes was more modest (Additional File 2: Table S3). In sum, these data reveal unexpected and dynamic interactions between Tau and aging and their divergent impact on the transcriptome and proteome.

### Tau-induced gene expression networks implicate aging and innate immune pathways

In order to reveal the broader biological processes disrupted by Tau, we next performed overrepresentation analysis using gene ontology (GO) annotations (Additional File 2: Table S5). We again focused on the Tau^R406W^ age-adjusted dataset, given the greater number of differential expression changes, and we initially examined the transcriptome. Complementary analyses of *Tau*^*WT*^ are included in the supplemental data. Among all differentially expressed transcripts, we detected significant enrichment for genes implicated in the cytoskeleton (p<1×10^−20^), endocytosis (p=8.9×10^−10^), synaptic function (p=1.7×10^−6^), innate immunity (p=8.9×10^−6^), and translation (p=8.4×10^−4^). One potential limitation of this approach is that it considers the entire transcriptome as a single regulatory unit and may therefore be underpowered to detect more restricted network modules. Therefore, in order to partition the transcriptomic data into coregulated gene sets, we implemented unsupervised hierarchical clustering and defined 6 discrete gene sets (n=33-1,863 genes; Additional File 2: Table S6), equally divided between Tau-associated up- and down-regulated groups (Fig. 3). As expected, each cluster was significantly enriched for genes corresponding to the biological pathways outlined above (Additional File 2: Table S7), consistent with identification of discrete transcriptional regulatory networks. Four gene clusters revealed strong age-dependent changes in both control and Tau flies, including both age-dependent decreases (clusters 1 & 3) or increases (clusters 2 & 4). As expected, these clusters (1-4) strongly overlap with age-associated gene expression changes obtained from controls (mean=78%, range 66-85% overlap). Interestingly however, we observe 2 distinct patterns for the relationship between Tau- and aging-associated transcriptome changes. First, in gene sets enriched for innate immune (cluster 2, increasing with age) or synaptic biology (cluster 1, decreasing with age), Tau amplifies the “aging expression signature”. Conversely, in clusters 3 and 4—enriched for endocytic and chromatin biology, respectively—Tau opposes the age-associated changes. Thus, these 2 alternate patterns conform to accelerated versus delayed brain aging, based on the transcriptome responses. In contrast, neither of the remaining clusters reveal strong age-dependent changes in control flies, with Tau triggering decreased (cluster 5) or increased (cluster 6) gene expression. Interestingly, in cluster 5, enriched for mitochondrial and epithelial gene sets, the Tau-induced downregulation in the transcriptome appears to be attenuated by aging. As a complementary strategy to define Tau-associated gene regulatory networks, we also implemented weighted gene correlation network analysis (WGCNA), identifying 15 mutually exclusive transcriptional modules (Additional File 2: Table S6). Among these, we found 7 modules significantly associated with Tau genotype in Tau^R406W^ flies (Additional File 1: Figure S7). Moreover, these modules substantially overlap with the gene sets defined using hierarchical clustering, resulting in similar functional enrichment profiles and recapitulating consistent interrelationships with aging (Additional File 2: Tables S6, S7).

**Figure 3:**
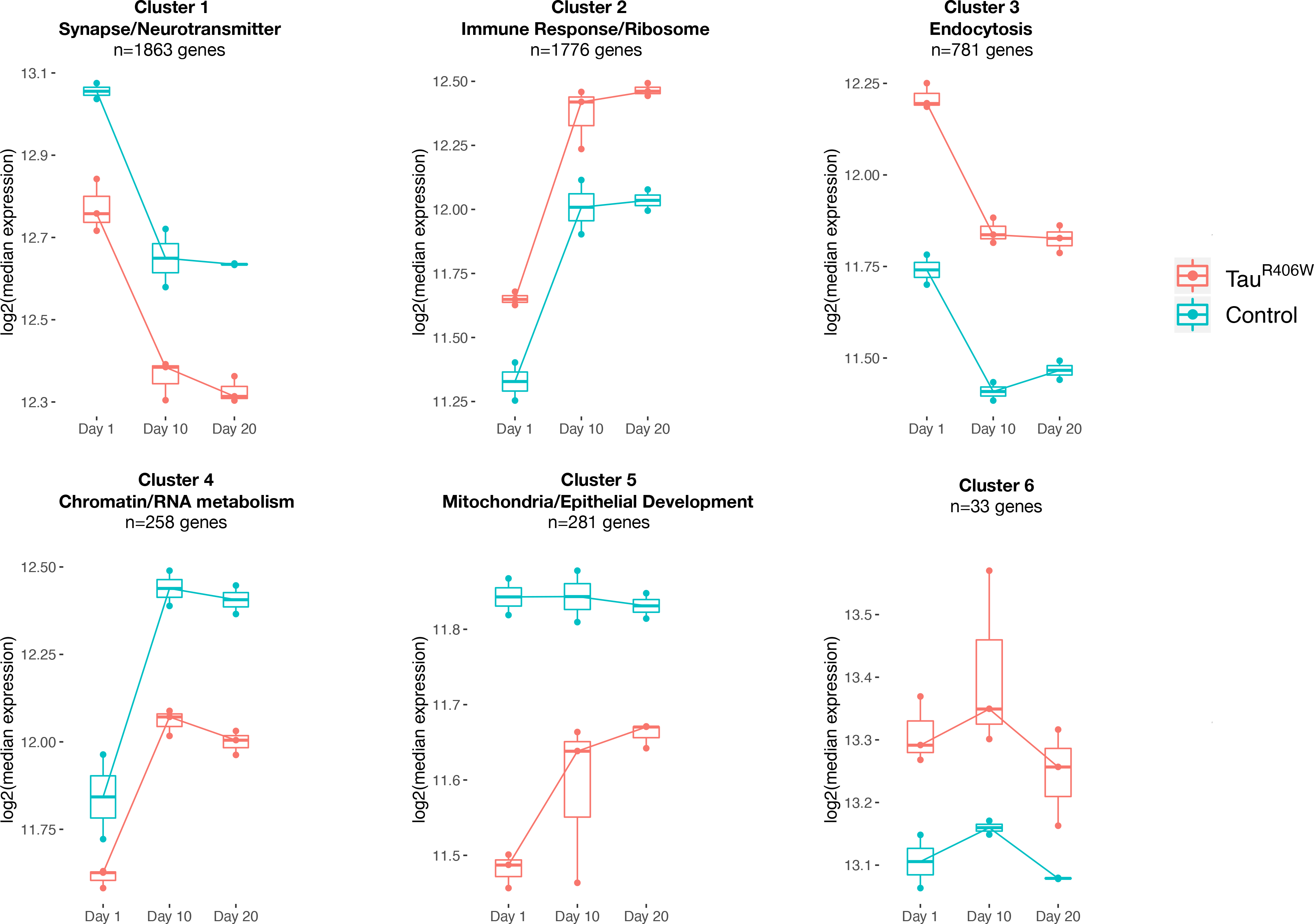
Tau-triggered gene expression clusters. Hierarchical clustering identified 6 gene sets with related Tau-induced expression patters (See also heatmap in Fig. 1a). Boxplots show log2-transformed median expression of genes within each cluster, including *elav>Tau*^*R406W*^ (Tau, red) and *elav* (Control, blue). Clusters are annotated based on size and significantly enriched gene ontology terms. See also Additional File 1: Figure S7, S8 and Additional File 2: Tables S6, S7.

In parallel analyses of the Tau^R406W^ proteome dataset, we detected enrichment among differentially expressed proteins for translation (p=1.2×10^−7^) and amino acid biosynthesis (p=7×10^−3^), including a preponderance of ribosomal proteins (p<1×10^−20^) (Additional File 2: Table S5). We next integrated the transcriptome derived clusters with complementary data from proteomics. Consistent with our analyses described above, we found variable concordance among the clusters, based on the direction of differential expression detected in the proteome (Additional File 1: Figure S8). For example, clusters enriched for immune and synaptic function were reciprocally up- or down-regulated in Tau animals, but both gene sets were predominantly (~60%) concordant in the proteome—differentially expressed proteins show consistent direction of change in Tau^R406W^ flies. By contrast, cluster 5, implicated in mitochondrial and epithelial biology, showed only 39% concordance, suggesting opposing regulatory interactions between the transcriptome and proteome.

### Cross-species annotation of Tau-specific changes from human brain gene expression profiles

Gene expression analysis from human postmortem brain tissue is confounded by mixed pathologies, making it difficult to identify those changes that are specifically triggered by Tau versus aging or other brain lesions. In contrast, our transcriptomic and proteomic analyses in flies benefit from matched experimental controls and longitudinal sampling, allowing definitive identification of Tau-triggered changes. We therefore leveraged our *Drosophila* results to annotate potential Tau-specific transcriptional changes from human brain gene expression profiles. We focused on 3 published analyses of differential gene expression, in relation to (i) AD clinical-pathologic diagnosis (n=478 cases / 300 controls; [10]), (ii) PSP clinical-pathologic diagnosis (n=82 cases / 76 controls; [9]), or (iii) neurofibrillary tangle pathologic burden (n=478 brains; [11]). As expected, following homology mapping using the *Drosophila* Integrated Ortholog Prediction Tool [43]; 57-66% of human genes had well-conserved fly homologs. The results of lookups are summarized in Table 3, and detailed results are included in Additional File 2: Table S8. In all 3 datasets, roughly half of conserved, differentially expressed changes are nominated as directly triggered by Tau pathology, based on cross-species annotation. Importantly, the observed human-fly overlaps appear more likely than that expected due to chance (hypergeometric test: AD, p=1.36×10^−63^; PSP, p=8.63×10^−19^; tangle burden, p=1.81×10^−48^). Moreover, ~50-60% of overlapping differentially expressed genes were concordant across species (i.e. gene up-regulation in both human AD and *Drosophila* Tau transcriptomes) (Additional File 2: Table S8). In a complementary analysis, we also examined the differentially expressed gene sets from human postmortem brains for overlaps with *Drosophila* aging-induced gene expression changes (Table 3 and Additional File 2: Table S8). An even greater proportion (~70%) of human genes altered in human tauopathy showed conserved changes during brain aging in flies. In fact, few human genes specifically overlapped with the Tau dataset, with 90% *overlapping both* the Tau- and aging differentially expressed gene sets. Lastly, we leveraged our fly proteomic data to annotate a recently reported mass-spectrometry dataset of differentially expressed proteins from 453 human brains, including 196 AD clinical-pathologic cases and 257 controls [19]. Despite the reduced depth of coverage for proteomics, this additional analysis highlights 63 proteins differentially expressed in human AD for which fly protein homologs are similarly dysregulated in response to Tau; 471 proteins overlapped with the complementary fly aging-dysregulated proteins.

**Table 3:**
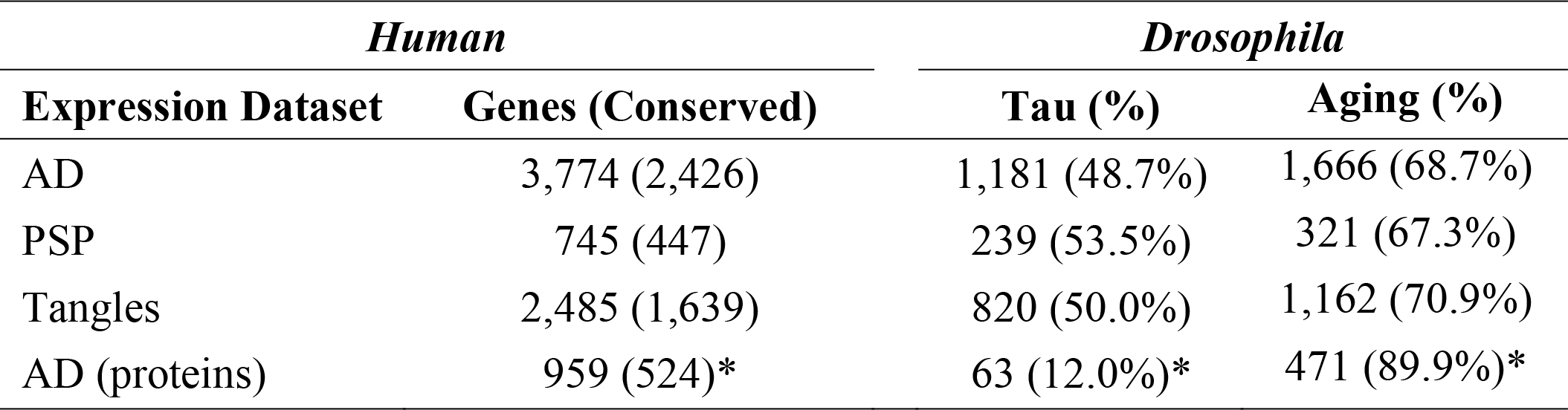
Tau- and aging-induced changes from cross-species overlaps. We examined differentially expressed transcripts from published RNA-seq analyses of human postmortem brain, including AD cases/controls [10], PSP cases/controls [9], or quantitative neurofibrillary tangle burden [11]. We also considered complementary mass-spectrometry proteomics from AD brains [19]. The total number of unique, differentially expressed human genes are noted along with the subset that are conserved in *Drosophila*. Among conserved genes, we examined the number and percentage with Tau- or aging-triggered differentially expressed homologs in flies. Given the reduced coverage of proteomics, we only consider conserved human proteins in which the homologous fly proteins were also detected in our assay. See Additional File 2: Tables S8 for complete results.

| *Human* |  | *Drosophila* |  |
| --- | --- | --- | --- |
| Expression Dataset | Genes (Conserved) | Tau (%) | Aging (%) |
| AD | 3,774 (2,426) | 1,181 (48.7%) | 1,666 (68.7%) |
| PSP | 745 (447) | 239 (53.5%) | 321 (67.3%) |
| Tangles | 2,485 (1,639) | 820 (50.0%) | 1,162 (70.9%) |
| AD (proteins) | 959 (524)* | 63 (12.0%)* | 471 (89.9%)* |

### Resolving amplifying versus protective expression changes using genetic modifiers

Tau-associated gene expression changes are excellent candidates for causal mechanisms in tauopathies—those with the potential to alter disease onset, progression, and/or neurodegeneration (Fig. 4). Alternatively, differentially expressed genes may define non-causal perturbations—such changes may represent candidates as biomarkers for the neuronal injury accompanying neurofibrillary tangle pathology. In order to identify potential causal gene expression changes, we integrated our findings with available results from 3 published, unbiased *Drosophila* screens, together defining 84 genetic modifiers of Tau-mediated neurotoxicity [25–27]. Among these, 37 genes were differentially expressed in the transcriptome and/or proteome (either Tau^WT^ or Tau^R406W^) (Table 4 and Additional File 2: Table S9). Next, for each of these 37 genes, we examined the direction of modifier tests from the literature to resolve whether the Tau-induced gene expression changes (up- or down-regulation) represent “amplifying” versus “protective” responses—that is, whether the observed perturbation in expression likely mediates or rather compensates for Tau-induced neuronal injury (Fig. 4). Up-regulated genes were defined as “amplifying” if genetic knockdown suppressed Tau toxicity and/or if overexpression reciprocally enhanced Tau phenotypes. For example, expression of *Ubiquitin activating enzyme 1* (*Uba1*), a regulator of axon pruning, autophagy, and apoptosis [44–46], is significantly increased in Tau^R406W^ flies. In published work [26], overexpression of *Uba1* enhanced Tau-induced retinal degeneration, suggesting that the observed up-regulation likely promotes (amplifies) Tau toxicity. Conversely, expression of *Mi-2*, encoding a CHD-family, chromatin-remodeling enzyme, is significantly decreased in *Tau*^*R406W*^ flies; however, since *Mi-2* is a loss-of-function suppressor of Tau neurotoxicity [27], we annotate this as a compensatory (protective) change. Interestingly, *Uba1* and *Mi-2* are members of expression clusters 2 & 4, respectively (Fig. 3 and Additional File 2: Table S6), which are similarly characterized by age-associated up-regulation but reveal opposing Tau-associated perturbations. Overall, we identify 18 amplifying (A) and 19 protective (P) gene expression changes induced by Tau. Thus, our transcriptome and proteome data can be integrated with genetic modifier studies to reconstruct a causal chain linking Tau, specific gene expression perturbations, and neurodegeneration. Moreover, 21 out of the 37 genes with published modifiers are also differentially expressed in one of the human datasets (Table 4 and Additional File 2: Table S8). For example, *CHD5* (homolog of *Mi-2*) is decreased in human brains with AD pathology. Based on the complementary studies of fly *Mi-2*, we can infer a potential Tau-triggered protective perturbation. These results demonstrate how *Drosophila* gene expression and genetic manipulation can be integrated to annotate human data for potential causal changes.

**Figure 4:**
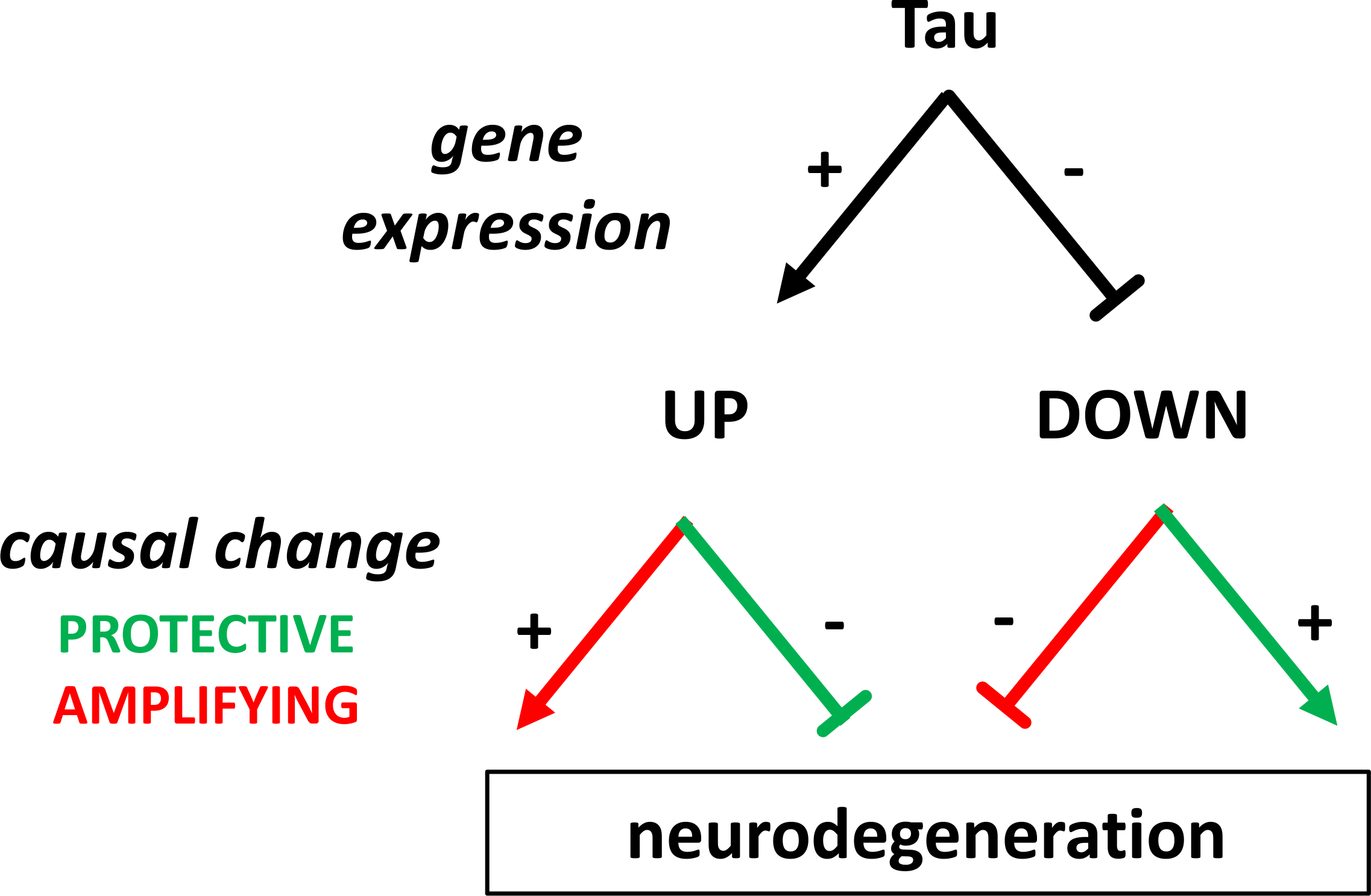
Model for integrating Tau-induced gene expression changes and modifiers. Schematic diagram illustrating the relationship between Tau-induced perturbations in gene expression and potential impact on neurodegeneration. Tau may cause up- or down-regulation for a given gene of interest, and either change may amplify (red) or protect against (green) neurotoxicity. Recapitulating the observed gene expression change through experimental manipulations and observing the consequences for neurodegenerative phenotypes permits reconstruction of the causal chain. See Table 4 for specific examples.

**Table 4:** Integration of gene expression with genetic modifiers. All *Drosophila* genes listed (at left) are modifiers of Tau-mediated neurodegeneration based on published unbiased screens [25–27]. Direction of Tau-induced differential expression is noted, including up- (top) or down-regulation (bottom) of transcripts. Based on the results of modifier tests, we can infer whether the observed Tau-induced expression perturbation is amplifying (A) or protective (P) for Tau neurotoxicity. See also Figure 4 and Additional File 2: Table S8 and S9. For each fly gene, we also note whether human gene homolog(s) are differentially expressed in human postmortem brain tissue from published analyses of AD [10] and neurofibrillary tangle burden [11]. In cases where the direction of expression was concordant with *Drosophila*, the human gene name is indicated in boldface. In a smaller PSP dataset [9], only 1 fly gene, *mub*, had a differentially-expressed human homolog, *PCBP4*.

| *Drosophila* |  | *Human* |  |
| --- | --- | --- | --- |
| Gene | Amplifying vs.<br>Protective | AD | Tangles |
| **<u>up-regulated</u>** |  |  |  |
| *cher* | A |  |  |
| *Uba1* | A |  |  |
| *Myd88* | A | *MYD88* |  |
| *CG10889* | A | *ZC3H12C* |  |
| *CG7970* | A |  |  |
| *Elf* | A | *GSPT1* | *EIF2S1* |
| *Fs(2)Ket* | A |  |  |
| *Mekk1* | A |  |  |
| *Nrg* | A | *CHL1* | *NRCAM* |
| *smid* | A |  |  |
| *Diap1* | A | *BIRC3* |  |
| *wun* | A | *PLPP1* |  |
| *Hop* | P | *STIP1, JAK1,*<br>*JAK2, JAK3* | *JAK3, TYK2* |
| *RpS21* | P | *RPS21* |  |
| *Past1* | P | *EHD2* |  |
| *Tis11* | P | *ZFP36L1,*<br>*ZFP36L2* |  |
| *w* | P |  |  |
| *Gbs-70E* | P | *PPP1R3C* |  |
| *cher* | P |  |  |
| *dally* | P |  |  |
| **<u>down-regulated</u>** |  |  |  |
| *fry* | A |  | *FRYL* |
| *Ptp4E* | A |  |  |
| *Atg6* | A |  |  |
| *Fmr1* | A |  |  |
| *mub* | A |  |  |
| *Bacc* | A |  |  |
| *jing* | P |  | *AEBP2* |
| *E(bx)* | P |  | *BPTF* |
| *tou* | P |  | *BAZ2B* |
| *jar* | P |  |  |
| *Mi-2* | P | *CHD5,*<br>*CHD4* | *CHD4* |
| *sgg* | P |  |  |
| *milt* | P |  | *TRAK2* |
| *stg* | P | *CDC25B* | *CDC25B* |
| *twe* | P | *CDC25B* | *CDC25B* |
| *Atx2* | P |  | *ATXN2L* |
| *CG7231* | P |  |  |

## Discussion

We present an integrated, longitudinal analysis of the aging brain in *Drosophila* tauopathy models. Our results identify perturbations affecting thousands of transcripts and hundreds of proteins triggered by expression of human Tau and highlight many promising biological pathways that likely contribute to neurodegeneration in tauopathy. Among these, regulators of innate immunity, the cytoskeleton, endocytosis, and synaptic transmission, have independent support from genome-wide association studies of AD [47, 48], neurofibrillary tangle burden [49], and PSP [50], consistent with causal roles. Although expressed at similar levels, Tau^R406W^, which causes familial FTD, was associated with a stronger impact on gene expression than Tau^WT^, inducing up to 7-fold increased response in gene expression. This result is consistent with the enhanced neurotoxicity of Tau^R406W^ in *Drosophila* [24, 51], as well as the more aggressive clinical profile of familial FTD compared to late-onset AD [1, 2]. Nevertheless, we observed strong overlap in the differential expression signatures of wildtype and mutant Tau, suggesting shared mechanisms.

Prior studies have profiled brain gene expression in Tau transgenic animals, including in flies [31, 32, 52] and mouse models [21, 23, 41]; however, none to our knowledge have longitudinally assessed both transcripts and proteins in parallel. Our analyses provide a glimpse of the dynamic regulatory crosstalk between the brain transcriptome and proteome that respond to brain injury, as in tauopathy. Remarkably, 42% of Tau-induced expression changes were discordant, with transcript and protein changing in opposite directions. This result is largely consistent with other emerging findings of surprisingly poor correlation between mRNA and protein levels among a variety of experimental systems [13, 17, 53–55], including analyses of human postmortem brain tissue [18, 19, 56]. In one notable study relevant to AD, a similar fraction (40%) of differentially expressed transcripts in the 5×FAD *amyloid precursor protein* transgenic mouse showed discordant changes in the proteome [57]. Many discordant changes likely reflect regulatory feedback interactions that maintain protein homeostasis. Consistent with this, we found that transcript-protein concordance varied among coexpressed, and therefore likely coregulated, gene sets. Our longitudinal data also provides clues to primary perturbations in selected cases (e.g. Sar1 in Figure 2); however, additional studies will be needed to confirm. Ultimately, successful translation from expression profiling studies requires unambiguous determination of whether a gene of interest is up- or down-regulated, but interpretation is currently limited by transcriptome-only analyses in most cases. Indeed, whether for nomination of potential therapeutic targets or development of diagnostic biomarkers, it will be essential to understand consequences at the protein level. We note that, among all differentially expressed transcripts, only a minority (~4%) were significantly and concordantly differentially expressed proteins--the remainder were either non-significantly differentially expressed, significant but discordant, or not detected at all by proteomics. Nevertheless, despite the comparatively reduced coverage of proteomics (~2,800 proteins vs. ~17,000 transcripts), many differentially expressed genes would not have been detected at all based on isolated transcriptional profiling. Thus, our results reinforce the value of proteomics for future investigations.

Age is the strongest known risk factor for AD, and aging-dependent progression is a defining feature of AD and other neurodegenerative tauopathies. As in studies of human postmortem tissue, most gene expression analyses of tauopathy models have been cross-sectional, partially obscuring the impact of aging and potential interactions with Tau-mediated changes. In our analyses, the majority (~70%) of Tau-triggered transcripts or proteins overlapped with those changes observed in aged control animals. Importantly, our longitudinal experimental design permitted identification of Tau-associated expression changes robust to aging adjustment. Remarkably however, even after adjustment, most Tau-mediated perturbations overlap with those seen in aging, and our cross-species analysis suggests consistent results for human tauopathy expression signatures. In short, our findings suggest that Tau pathology primarily modulates the endogenous gene expression programs of brain aging. Indeed, following hierarchical clustering, 4 out of 6 differentially expressed gene sets mirrored aging expression patterns, consistent with either Tau-accelerated or delayed aging. These complementary patterns may represent disease amplifying or protective responses, respectively, as shown for *Uba1* and *Mi-2*. Interestingly, aging was associated with a quantitatively enhanced transcriptional signature in the Tau transgenic animals, characterized by an 18% increase in differentially-expressed genes. Reciprocally, Tau expression was accompanied by a 78% reduction in age-associated changes in the proteome. Though further investigation is warranted to confirm these observations, our analyses define Tau expression signatures in the both transcriptome and proteome enriched for genes implicated in translation, including numerous ribosomal proteins. Consistent with this, pathologic forms of Tau have been shown to avidly bind ribosomal proteins and disrupt their function [40–42].

While aging has myriad systemic and cellular targets, one key emerging theme is the dysregulation of innate immune mechanisms leading to a systemic pro-inflammatory state, which has been termed “immunosenescence” or “inflamm-ageing” [58, 59]. In our analysis, innate immune pathways were strongly enriched among both aging- and Tau-associated, differentially expressed genes, and this result is consistent with brain gene expression profiling in mouse models of healthy aging [23, 60] and tauopathy [61, 62]. Similarly, multiple transcriptome- and proteome-wide analyses of human postmortem brain from AD or other tauopathies, such as PSP, have identified evidence of dysregulated immune pathways [9–11, 18], and similar signatures have been implicated in brains from aged individuals without known neurodegenerative disease [63, 64]. Importantly, genome-wide association studies in AD highlight an abundance of susceptibility gene candidates implicated in immune regulation (e.g. *TREM2, CD33, CR1*), strongly suggesting a causal role in disease pathogenesis [48]. Further, polygenic modeling [65] and analyses of human cortical transcriptomes [66] converge to implicate activated microglia in the development of Tau pathology and susceptibility for AD. Numerous follow-up studies, including in mouse and cellular models, implicate microglia and astroglia with potential roles in propagating a pathogenic inflammatory cascade [67]. However, the prevailing mechanistic models of neuroinflammation in AD have largely focused on amyloid-beta as an upstream trigger and tau pathology as a downstream consequence, and the role of aging *per se* is often minimized. Nevertheless, primary tauopathies lacking amyloid pathology, such as PSP, and corresponding mouse models manifest prominent neuroinflammatory brain expression signatures. By contrast with mammals, neurons significantly outnumber glia in the *Drosophila* brain, and true microglial cells are not present in invertebrates [68]. Nevertheless, innate immune pathways are evolutionarily ancient, and toll-like receptor signaling components are not only expressed in fly neurons and glia, but they are required for brain maintenance in aging [39, 69]. In the future, single-cell RNA-seq in *Drosophila* models of tauopathy may permit dissection of which cell types generate immune expression signatures along with complementary cell-type specific manipulations to confirm potential causal roles.

Gene expression profiling has emerged as a promising tool for functional genomic dissection of AD and other tauopathies; however, interpretation of these data can be powerfully enhanced by integration with complementary studies in model organisms. We have performed several cross-species analyses to highlight applications of our *Drosophila* tauopathy resource. One important challenge is to differentiate those gene expression changes specifically provoked by Tau-mediated mechanisms. Besides the influence of aging and life experiences, human brains commonly accumulate mixed pathologies [16]. By contrast, experimental models permit precisely-controlled manipulations that can isolate the responsible causal triggers. Roughly half of all conserved, differentially-expressed genes from the largest available analyses of human AD or PSP brain tissue were annotated as Tau-induced perturbations based on our *Drosophila* experiment. Remarkably, an even larger proportion of expression changes (70%) were triggered by aging and we observed virtually complete overlap between Tau- and aging-associated
changes. This result reinforces the intimate connection between the impact of neurodegenerative pathologies and aging on brain gene expression. Another major challenge following human gene expression analyses is to differentiate proximal causal pathways from more downstream, non-causal consequences of neurodegeneration. Experimental models permit controlled manipulations that mimic observed expression changes along with assessments to define potential impact on neurodegenerative phenotypes. In particular, *Drosophila* offers high-throughput genetics enabling unbiased, large-scale genetic screens for modifiers of Tau-mediated neurotoxicity [25–27]. By integrating these results with our RNA-seq findings, and cross-referencing with human gene expression profiles, we successfully highlight genes altered in human tauopathy that are strong candidates for further investigation as either amplifying or protective causal modifiers. In the future, targeted genetic manipulations of other conserved, differentially-expressed transcripts and/or proteins will significantly extend the value of our cross-species resource.

## Methods

### Drosophila stocks and husbandry

*UAS-Tau*^*WT*^ and *UAS-Tau*^*R406W*^ transgenic flies, as previously described in Wittmann et. al. 2001, were crossed with the pan-neuronal expression driver *elav-GAL4* to generate experimental animals with the genotype *elav-GAL4/+;UAS-Tau*^*WT*^/+ or *elav-GAL4/Y;UAS-Tau*^*WT*^/+ (*elav > Tau*^*WT*^) and *elav-GAL4/+;UAS-Tau*^*R406W*^/+ or *elav-GAL4/Y;UAS-Tau*^*R406W*^/+ (*elav > Tau*^*R406W*^), respectively. These flies express the human Tau 0N4R isoform (383 amino acids). For control animals, we used the genotypes: *elav-GAL4/+* and *elav-GAL4/Y*. All flies were raised on standard molasses-based *Drosophila* media at 25°C with ambient light conditions, and aged to 1-, 10-, or 20-days following eclosion. We confirmed expression of Tau at similar levels in *elav > Tau*^*WT*^ and *elav > Tau*^*R406W*^ flies using western blot analysis, as previously described [40] using the following antibodies: rabbit anti-Tau (1:5000, Dako); rabbit anti-GAPDH (1:5000, GeneTex) and HRP-conjugated anti-rabbit (1:10000, Santa Cruz).

### *Drosophila* RNA-sequencing data

The *Drosophila* RNA-sequencing (RNA-seq) dataset analyzed for this work was generated as part of another study, where it is described in detail [40]. Briefly, for *elav*>*Tau* and *elav* controls, animals were evaluated at 1-, 10-, or 20-days. To avoid possible batch effects, experimental and control genotypes used for each comparison (*Tau*^*WT*^ and *Tau*^*R406W*^) were sequenced together, such that 2 separate control datasets were generated (control 1 and control 2, respectively) for the Tau^WT^ and Tau^R406W^ RNA-seq analyses. Triplicate samples (n=3) were used for all genotypes and time points, except for the *elav* control genotype used for the comparison with *Tau*^*R406W*^(control 2), for which duplicate samples were used (n=2):

| <u>Exp</u> | <u>Replicates (n)</u> | <u>Ctrl</u> | <u>Replicates (n)</u> | <u>Age (d)</u> |
| --- | --- | --- | --- | --- |
| *Tau*<sup>WT</sup> | 3 | *elav* | 3 | 1, 10, 20 |
| *Tau*<sup>R406W</sup> | 3 | *elav* | 2 | 1,10, 20 |

Total RNA was extracted from approximately 100 adult fly heads (for each genotype/age/sample), equally divided between males and females. Sequencing was performed on the Illumina HiSeq with 100bp paired-end reads. Gene expression values from each sample were quantified as the number of reads mapped (to a specific gene) by setting --quantMode to GeneCounts in STAR 2.5.3a [70]. Unsupervised clustering of samples was assessed by UMAP using DESeq2 depth normalized read counts as described in [71].

### Protein extraction and mass-spectrometry

For proteomics, the identical genotypes (*elav>Tau*^*WT*^, *elav>Tau*^*R406W*^, and *elav*), time points (1-, 10-, or 20-days), and conditions were evaluated as for the RNA-seq analyses. Triplicate samples (n=3) were used for all genotypes and timepoints, and a single control series (control 3) was used since all samples were processed together. *Drosophila* proteomics was performed according to previously published protocols [18]. Each replicate (40 fly heads of 1:1 male/female ratio per sample) was homogenized in 500 uL of urea lysis buffer (8M urea, 100 mM NaHPO_4_, pH 8.5), including 5 μL (100× stock) HALT protease and phosphatase inhibitor cocktail (Pierce). Protein supernatants were transferred to 1.5 mL Eppendorf tubes and sonicated (Sonic Dismembrator, Fisher Scientific) 3 times for 5 s with 15 s intervals of rest at 30% amplitude to disrupt nucleic acids and subsequently vortexed. Protein concentration was determined by the bicinchoninic acid (BCA) method, and samples were frozen in aliquots at −80°C. Each brain homogenate was analyzed by SDS-PAGE to assess for protein integrity. Protein homogenates (150 μg) were diluted with 50 mM NH_4_HCO_3_ to a final concentration of less than 2M urea and then treated with 1 mM dithiothreitol (DTT) at 25°C for 30 minutes, followed by 5 mM iodoacetimide (IAA) at 25°C for 30 minutes in the dark. Protein was digested with 1:100 (w/w) lysyl endopeptidase (Wako) at 25°C for 2 hours and further digested overnight with 1:50 (w/w) trypsin (Promega) at 25°C. Resulting peptides were desalted with a Sep-Pak C18 column (Waters), dried under vacuum, and 2 μg was resuspended in peptide loading buffer (0.1% formic acid, 0.03% trifluoroacetic acid, 1% acetonitrile). Peptide mixtures were separated on a self-packed C18 (1.9 μm Dr. Maisch, Germany) fused silica column (25 cm × 75 μM internal diameter (ID); New Objective, Woburn, MA) by a NanoAcquity UHPLC (Waters, Milford, FA) and monitored on a Q-Exactive Plus mass spectrometer (ThermoFisher Scientific, San Jose, CA). Elution was performed over a 120-minute gradient at a rate of 400nL/min with buffer B ranging from 3% to 80% (buffer A: 0.1% formic acid and 5% DMSO in water, buffer B: 0.1% formic and 5% DMSO in acetonitrile). The mass spectrometer cycle was programmed to collect one full MS scan followed by 10 data dependent MS/MS scans. The MS scans (300-1800 m/z range,1,000,000 AGC, 150 ms maximum ion time) were collected at a resolution of 70,000 at m/z 200 in profile mode and the MS/MS spectra (2 m/z isolation width, 25% collision energy, 100,000 AGC target, 50 ms maximum ion time) were acquired at a resolution of 17,500 at m/z 200. Dynamic exclusion was set to exclude previous sequenced precursor ions for 30 seconds within a 10 ppm window. Precursor ions with +1, and +6 or higher charge states were excluded from sequencing.

Raw data for all samples was analyzed using MaxQuant v1.5.2.8 with Thermo Foundation 2.0 for file reading capability. The search engine Andromeda, integrated into MaxQuant, was used to build and search a Uniprot fly database consisting of 13704 target sequences, plus 245 contaminant proteins from the common repository of adventitious proteins (cRAP) built into MaxQuant. Methionine oxidation (+15.9949 Da), asparagine and glutamine deamidation (+0.9840 Da), and protein N-terminal acetylation (+42.0106 Da) were variable modifications (up to 5 allowed per peptide); cysteine was assigned a fixed carbamidomethyl modification (+57.0215 Da). Only fully tryptic peptides were considered with up to 2 miscleavages in the database search. A precursor mass tolerance of ±20 ppm was applied prior to mass accuracy calibration and ±4.5 ppm after internal MaxQuant calibration. Other search settings included a maximum peptide mass of 6,000 Da, a minimum peptide length of 6 residues, 0.05 Da tolerance for high resolution MS/MS scans. Co-fragmented peptide search was enabled to deconvolute multiplex spectra. The false discovery rate (FDR) for peptide spectral matches, proteins, and site decoy fraction were all set to 1 percent. Quantification settings were as follows: re-quantify with a second peak finding attempt after protein identification has completed; match MS1 peaks between runs; a 0.7 min retention time match window was used after an alignment function was found with a 20-minute RT search space. The quantitation method only considered razor plus unique peptides for protein level quantitation. Quantitation of proteins was performed using LFQ (label-free quantification) intensities given by MaxQuant. The full list of parameters used for MaxQuant are available as parameters.txt accompanying the public release (see Data Availability). Unsupervised clustering of samples with UMAP was performed using DEseq2 depth normalized LFQ values (Additional File 1: Figure S3). Missing proteomic LFQ values were imputed on a per sample basis as previously described in [72]. Missing values were imputed by drawing from a Gaussian distribution simulating expression near the LFQ detection limit, a down-shift of 1.8 standard deviations from the median sample expression. For quality assurance, we tabulated for each protein the number of replicate samples with complete data (non-imputed), broken down by genotype and age (Additional File 2: Table S10). For subsequent differential expression analyses, LFQs for the minority of proteins with multiple detected isoforms (n=83) were collapsed to a single value by taking the per sample mean. Abundance data for UniProt peptide IDs that did not map to a fly gene symbol were excluded from analysis.

### Analysis of differentially expressed transcripts and proteins

Differential-expression analysis of transcripts and proteins was performed using DESeq2 [73]. As detailed above, for transcriptome analyses, *elav>Tau*^*WT*^ or *elav>Tau*^*R406W*^ were compared with the batch-matched *elav* control data (control set 1 or 2, respectively). Genes with an average read count <50 across all samples in the comparison were excluded. For proteomic analyses, the single *elav* control set (control 3) was compared to either *elav>Tau*^*WT*^ or *elav>Tau*^*R406W*^, and absolute peptide counts (LFQ) were used as the input for DESeq2 (which only accepts integers). Raw transcript or peptide counts were normalized for library depth using DESeq2 median of ratios, and tested for differential expression using a generalized linear model. We initially determined Tau-induced differentially expressed transcripts or proteins cross-sectionally, examining Tau and control data separately at each time point (expression ~ genotype, stratified by age for either 1-, 10-, or 20-day old animals). Subsequently, we performed joint regression analyses incorporating all longitudinal data, and including a covariate for age (expression ~ genotype + age); the genotype term coefficient was used for significance testing. Genes and proteins in Figure 2 and Additional File 1: Figure S5 were plotted using log-transformed and depth-normalized expression or LFQ values. For determination of age-related changes in transcripts or proteins, our data was stratified by genotype, evaluating *elav* controls or *elav>Tau* flies separately, and age was used as the predictor variable. Differential expression was computed for either (i.) day 1 vs. day 10, (ii.) day 10 vs. day 20, or (iii.) day 1 vs. day 20. Significance testing was performed using the Wald test, implemented within DESeq2. In order to account for multiple-comparisons, the Benjamini-Hochberg procedure was applied, and a false discovery rate (FDR) < 0.05 was considered significant.

For analysis of concordance between transcriptome and proteome, we examined the sign (positive or negative) of the genotype coefficient from the longitudinal (joint) regression model. Concordant transcripts were defined as having consistent direction of change (e.g. either positive or negative fold-change). Functional enrichment for differentially expressed transcripts or proteins (joint regression model) was evaluated using the over-representation analysis (ORA) function of the WEB-based GEne SeT AnaLysis Toolkit [74]. All ORA analyses were conducted using the R implementation of WEBGESTALT. The minimum number of genes per category was set to 5. We employed the following databases: GO biological processes, GO molecular functions, GO cellular component, KEGG, and Panther. Enrichment significance was defined using Fisher’s exact test, followed by the Benjamini-Hochberg procedure; significance was set at FDR < 0.05.

### Hierarchical Clustering and WGCNA analysis

Hierarchical clustering was performed to evaluate Tau^R406W^-associated, differentially-expressed transcripts (n = 4,992 genes), based on the joint regression model (Fig. 1b). Normalized expression counts for differentially-expressed genes were used as input. Pearson correlation was used as the distance metric and the complete linkage was used for distance calculation. Heatmaps of hierarchically clustered transcripts were generated using the heatmap.2 function from the gplots package in R. Based on a non-negative matrix factorization (NMF) rank survey using the NMF package in R [75], the optimal number of clusters was determined to be 6, maximizing cophenetic scores while minimizing residuals (Additional File 1: Figure S9). This was applied to the clustering as a manual tree cut to yield 6 final clusters. Functional enrichment for cluster gene set was performed as described above. Concordance between transcripts in each cluster with corresponding proteins detected in the Tau^R406W^ proteomic data was further evaluated by comparing the directions of log2 fold-changes. Median expression counts of genes belonging to each cluster were calculated from normalized expression values from all replicates in each genotype (Tau^R406W^ or control set 2) and age.

Weighted gene coexpression network analysis (WGCNA) [76] was performed on expression counts from all *Tau*^*R406W*^ transcripts (n = 10,217 genes) after normalization in DESeq2 (median-of-ratios depth normalization). The soft threshold parameter was set at 5, deepSplit = 4, and minimum module size = 23. Expression behavior of WGCNA modules were summarized by calculating module “eigengenes”. Module eigengene is defined by PC1 loadings of a given module. Closely related modules were merged based on module eigengenes at a distance threshold of MEDissThres = 0.1. The cluster dendrogram and module membership of transcripts are displayed in Additional File 1: Figure S10. Module eigengenes of each of the 15 resulting modules was examined for correlation with the Tau^R406W^ genotype via Pearson correlation (Additional File 1: Figure S7). Normalized expression of genes in modules with module eigengenes that have significant correlation to the Tau^R406W^ genotype were further evaluated in Tau^R406W^ animals and controls (control set 2).

### *Drosophila* and human gene set overlaps

In order to evaluate human-fly gene set overlaps, we first determined the fly homologs for all human AD, tangle, or PSP differentially-expressed genes using the DRSC Integrated Ortholog Prediction Tool (DIOPT; [43]), applying a minimum DIOPT score threshold of 5 (Additional File 2: Table S8). Where more than one fly homolog had a DIOPT score > 5, all were included. We then computed enrichments of each human-derived data set (fly homologs) for either (i) Tau- or (ii) age-induced differentially expressed gene sets, based on our experimental analyses in *Drosophila* models, using the phyper base function of R to conduct a hypergeometric test. For Tau-induced fly genes, we include significant, differentially expressed genes from either the *Tau*^*R406W*^ or *Tau*^*WT*^ joint regression model (age-adjusted). For aging-induced fly genes, we considered all unique differentially expressed genes based on our analyses of *elav* control flies from multiple timepoints (1 vs. 10 days, 10 vs. 20 days, and 1 vs. 20 days), including from control sets 1 & 2 for transcriptome studies or the complementary proteomic control set. Input values for human-to-fly hypergeometric tests were:

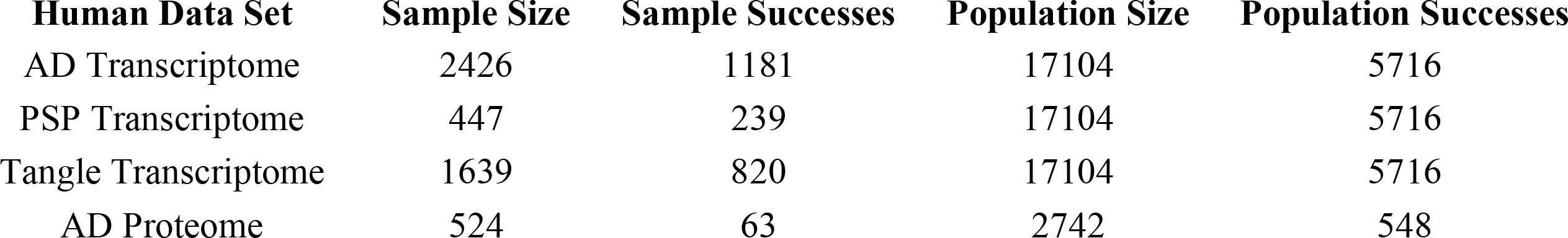

| Human Data Set | Sample Size | Sample Successes | Population Size | Population Successes |
| --- | --- | --- | --- | --- |
| AD Transcriptome | 2426 | 1181 | 17104 | 5716 |
| PSP Transcriptome | 447 | 239 | 17104 | 5716 |
| Tangle Transcriptome | 1639 | 820 | 17104 | 5716 |
| AD Proteome | 524 | 63 | 2742 | 548 |

For the analyses integrating *Drosophila* RNA-seq and published modifiers, we again used DIOPT to determine the human homologs of relevant fly genes (DIOPT score > 5).

## Declarations

### Ethics approval and consent to participate

Not applicable.

### Consent for publication

Not applicable.

### Availability of data and materials

The complete RNAseq and proteomics data from *Drosophila* used in this study are available for download from the AMP-AD Knowledge Portal (www.synapse.org/ampad) (doi: 10.7303/syn7274101). Detailed results of our analyses are also included with this article and its additional files.

### Authors’ Contributions

Conceptualization, C.G.M., T.W., H.K.Y., N.T.S., Z.L., J.M.S.; Investigation and Analysis, C.G.M., T.W.H.K.Y., Y.-C.H., C.G., E.B.D., D.D.; Writing—Original Draft, C.G.M., T.W., H.K.Y., J.M.S.; Writing— Review & Editing, C.G.M., T.W., H.K.Y., Y.-C.H., C.G., D.D., E.B.D, P.L.D., N.T.S., J.M.S., Z.L.; Funding Acquisition, P.L.D., N.T.S., J.M.S., Z.L.; Supervision, N.T.S, J.M.S., Z.L.

## Acknowledgements

We thank M.B. Feany for generously sharing *Drosophila* stocks.

