## Additional File 1 for "Integrated analysis of the aging brain transcriptome and proteome in tauopathy"

**ADDITIONAL FILE 1: SUPPLEMENTAL FIGURES**

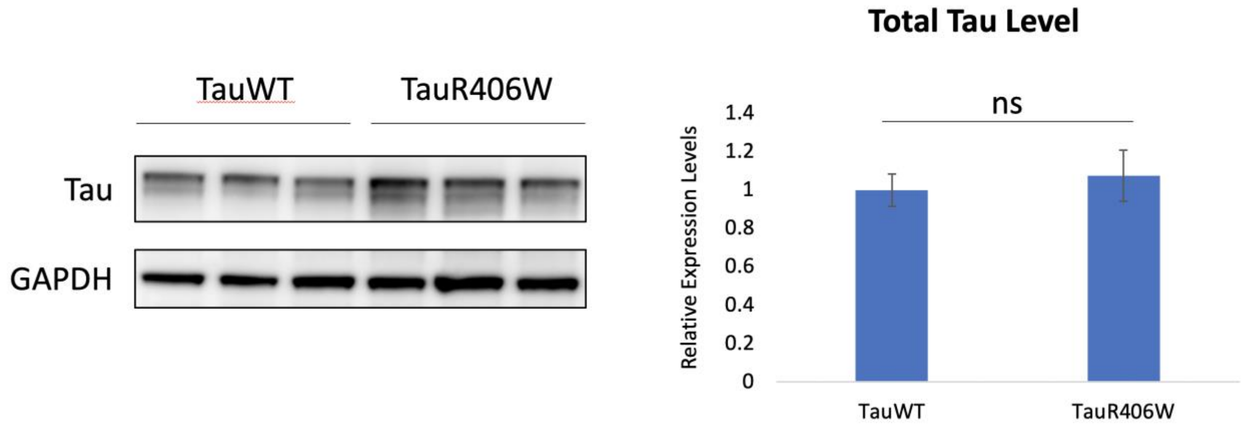


**Figure S1.** Human Tau protein is expressed at comparable levels in *elav>Tau$_{WT}$* and *elav>Tau$_{R406W}$* transgenic flies. Western blot quantification of Tau protein expression from 1-day-old animals (n=3 per genotype). Statistical analysis was performed using student's t-test.

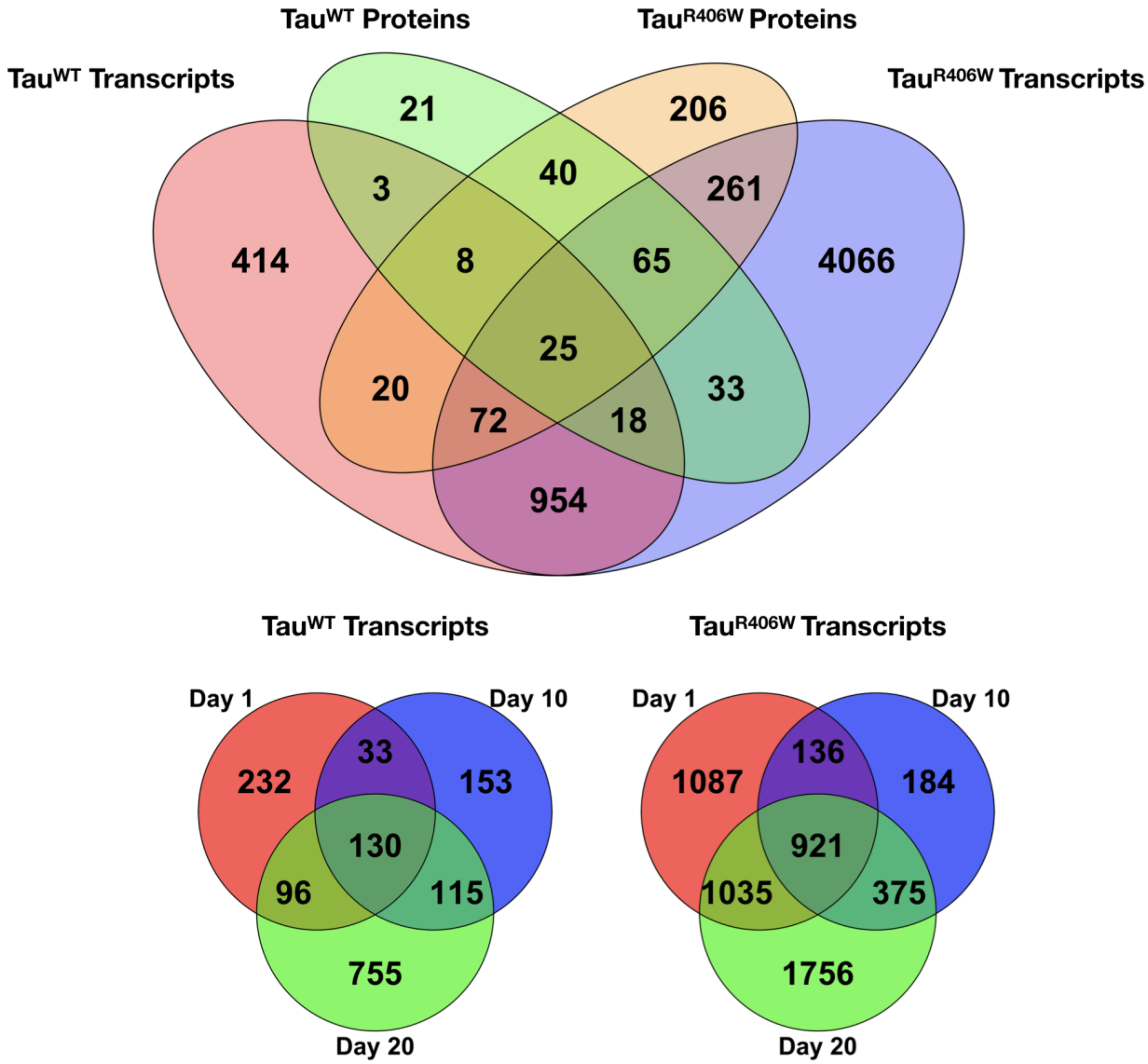


**Figure S2. (**Top**)** Venn diagram showing the number of differentially expressed genes shared between the transcriptome and proteomes of either *elav>Tau$_{WT}$* or *elav>Tau$_{R406W}$* transgenic flies, when compared with *elav* controls. Results of cross-sectional analyses stratified by age were combined (total of 1-, 10-, and 20-days). (Bottom) Venn diagrams showing overlaps of TauWT- (left) or TauR406W-induced differentially-expressed genes across the 3 ages evaluated (1-, 10-, and 20-days). See also Table 1 and Additional File 2: Table S1.

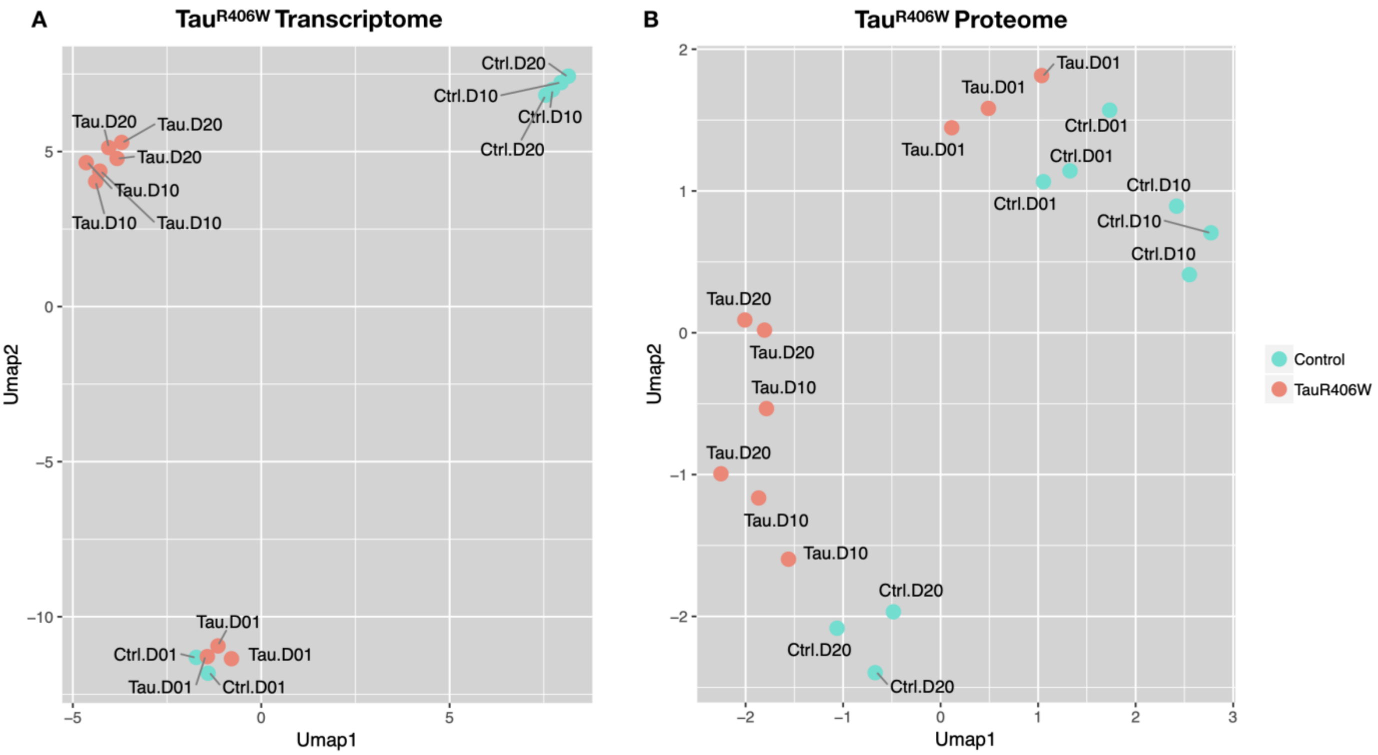


**Figure S3.** Age and Tau are major drivers of gene expression differences among samples. Uniform manifold approximation and projection (UMAP) of Tau$_{R406W}$ transcriptome (A) and proteome (B) expression data. Raw counts were normalized for library size and log-transformed. *elav* control (Ctrl, blue) and *elav>Tau$_{R406W}$* (Tau, red) samples are shown, including from 1-, 10-, and 20-day old animals (D01, D10, D20, respectively). See also Fig. 1a, b.

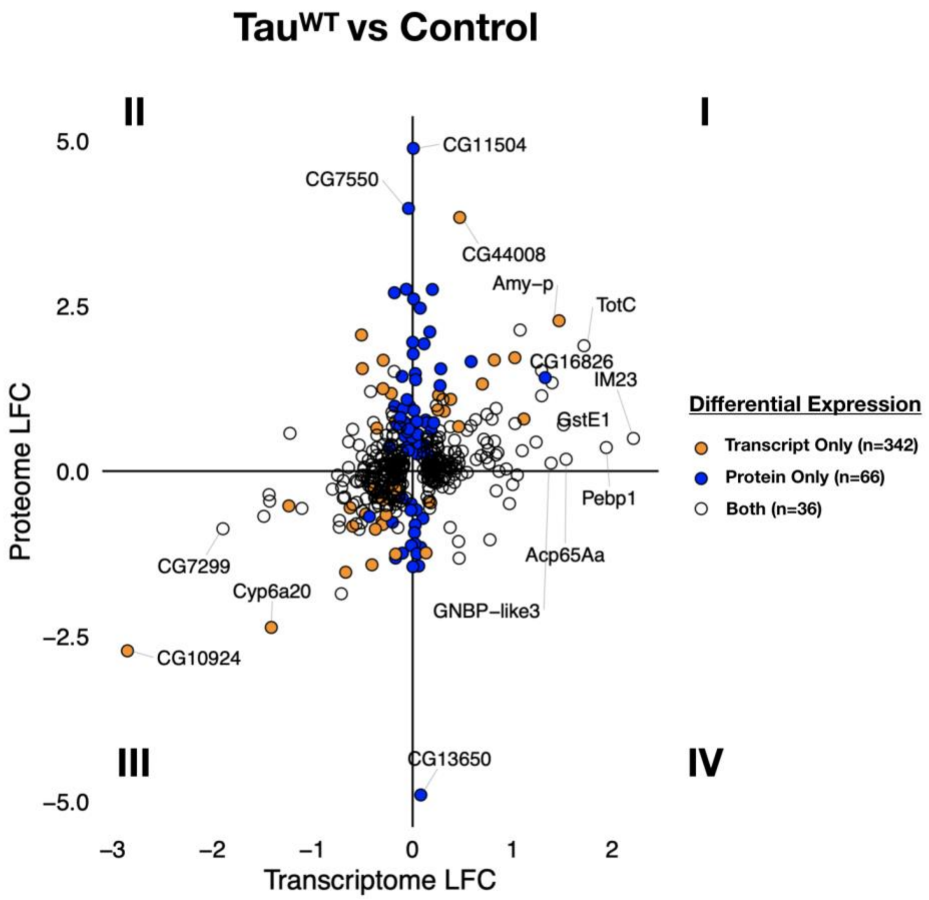


| Differential Expression | Concordant (I + III) | Discordant (II + IV) |
|---|---|---|
| Transcripts | 251 (66%) | 127 (34%) |
| Proteins | 63 (62%) | 39 (38%) |

**Figure S4.** Plot (top) showing $Tau_{WT}$-triggered $log_2$ fold-change (LFC) in the transcriptome and proteome. The plot only includes those genes detected as both transcripts and proteins and also differentially expressed (n=446, FDR < 0.05), based on the joint regression model including longitudinal data and adjusting for age. Colors denote whether gene was differentially expressed in the transcriptome (unfilled), proteome (blue), or both (orange). Quadrants I and III include gene expression changes that are concordant (same direction) at the transcript and protein level; whereas quadrants II and IV depict discordant changes. A substantial proportion of differentially-expressed transcripts or proteins are discordant (table, bottom).

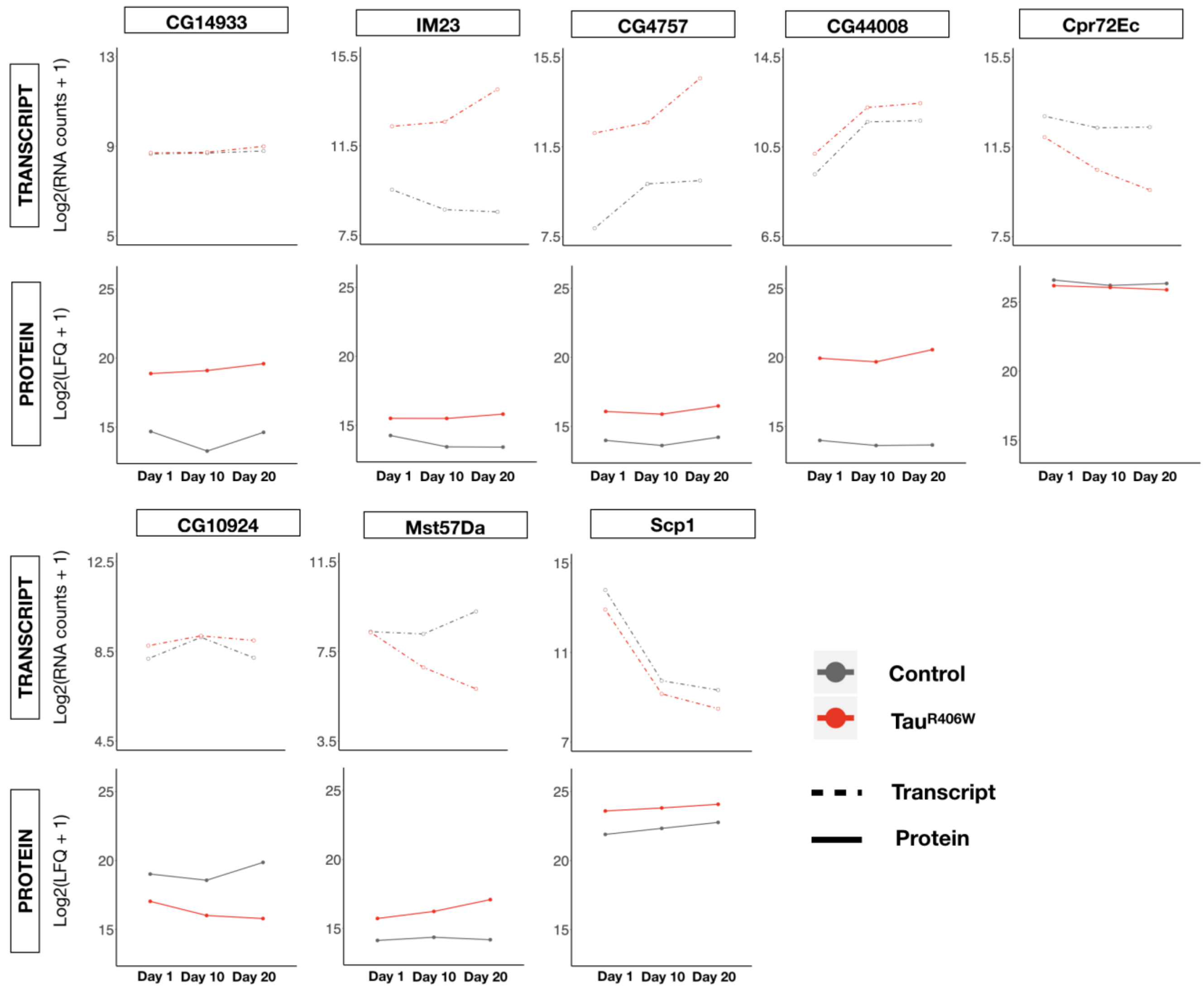


**Figure S5.** log2-transformed expression of selected genes in *elav>Tau$_{R406W}$* (Tau, red) and *elav* (Control, gray) is shown for transcriptome (depth normalized counts) and proteomes (normalized label-free quantification intensity (LFQ)). The selected genes are also noted in Fig. 1c. and are differentially expressed (FDR < 0.05) in both the transcriptome and proteome, except CG14933 and CG10924 which are protein-only, based on the joint regression model including all longitudinal data and adjusting for age.

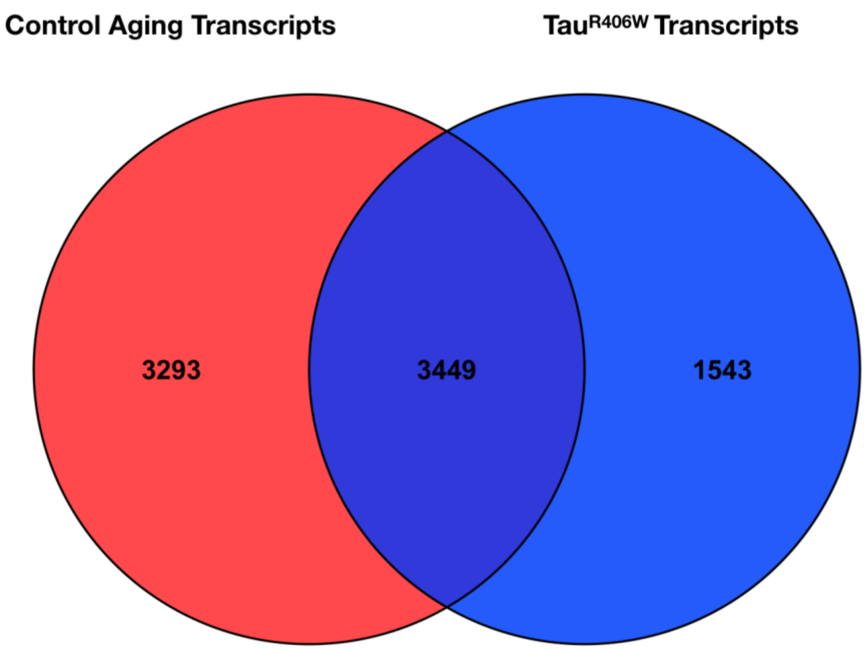


**Figure S6.** Venn diagram showing overlap between *elav*>*Tau$_{R406W}$*-induced (blue) differentially-expressed transcripts and aging-induced changes (red) detected from batch-matched *elav* controls. Tau$_{R406W}$ -induced differential expression is based on the joint regression model including all longitudinal data and adjusting for age. The total number of aging-associated, differentially-expressed transcripts are based on the union of 3 comparisons (1- vs. 10-days, 10- vs. 20-days, and 1- vs. 20-days). See also Table 2 and Additional File 2: Tables S2, S3.

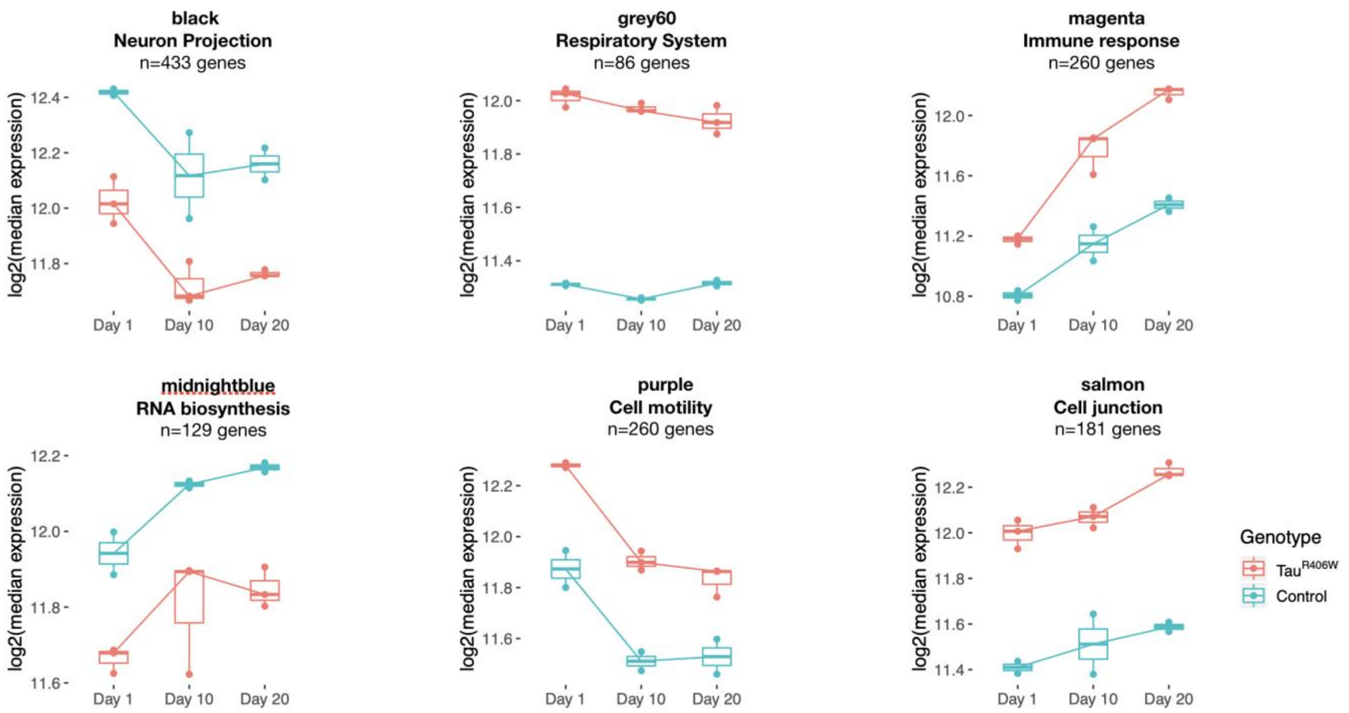


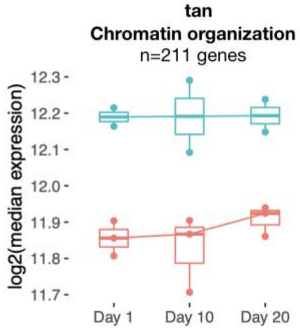


**Correlation to Tau$^{R406W}$**

| WGCNA module | Pearson R | pvalue |
|---|---|---|
| tan | -0.964 | $7.37 \times 10^{-9}$ |
| salmon | 0.926 | $7.33 \times 10^{-7}$ |
| grey60 | 0.961 | $1.28 \times 10^{-8}$ |
| magenta | 0.594 | $1.96 \times 10^{-2}$ |
| purple | 0.669 | $6.34 \times 10^{-3}$ |
| black | -0.83 | $1.27 \times 10^{-4}$ |
| midnightblue | -0.836 | $1.05 \times 10^{-4}$ |

**Figure S7.** Seven out of 15 WGCNA modules were significantly correlated with *elav*>*Tau*$_{R406W}$ genotype as described in the methods. Boxplots show log2-transformed median expression of genes within each of these correlated modules, including *elav*>*Tau*$_{R406W}$ (Tau, red) and *elav* (Control, blue). Clusters are annotated based on size and significantly enriched gene ontology terms. Pearson coefficients and p-values for module correlations to Tau$_{R406W}$ genotype are also shown. See also Additional File 2: Tables S6, S7.

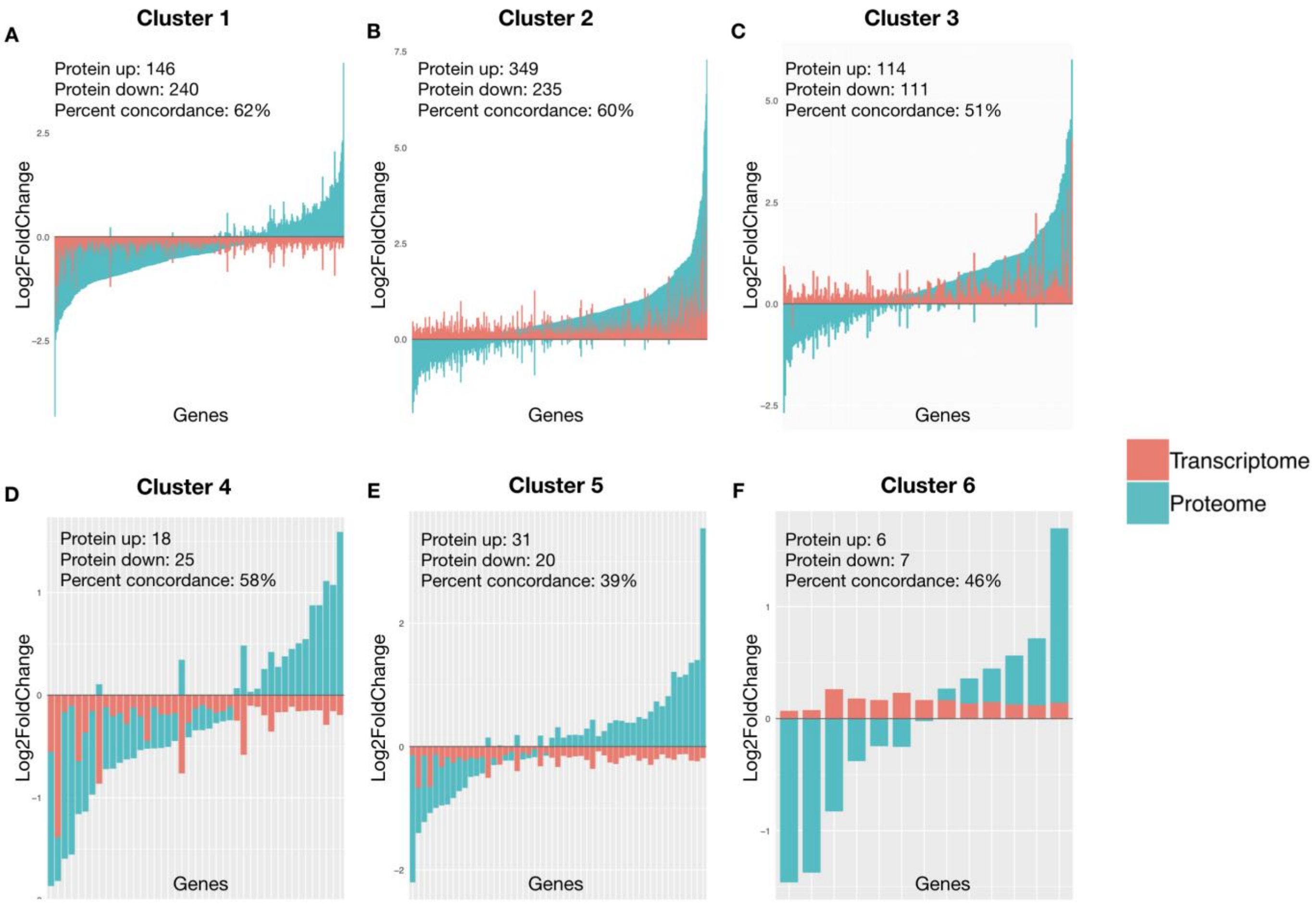


**Figure S8.** Integration of Tau-induced transcriptome and proteome changes in gene expression clusters. Unsupervised hierarchical clustering was performed on 4,992 differentially expressed transcripts (FDR < 0.05) in *elav>Tau$_{R406W}$* vs. *elav* controls, based on the joint regression model, including all longitudinal data and adjusting for age. Log2 fold-change of for gene members of the 6 resulting, mutually-exclusive clusters are shown. Stacked bar plots are restricted to those genes detected in both the transcriptome (red) and proteome (blue). The number of proteins that are either up- or down-regulated and the percent concordance between proteins and transcripts are noted. As a result of hierarchical clustering, transcripts within a given cluster have consistent direction of change (up or down). See also Fig. 3.

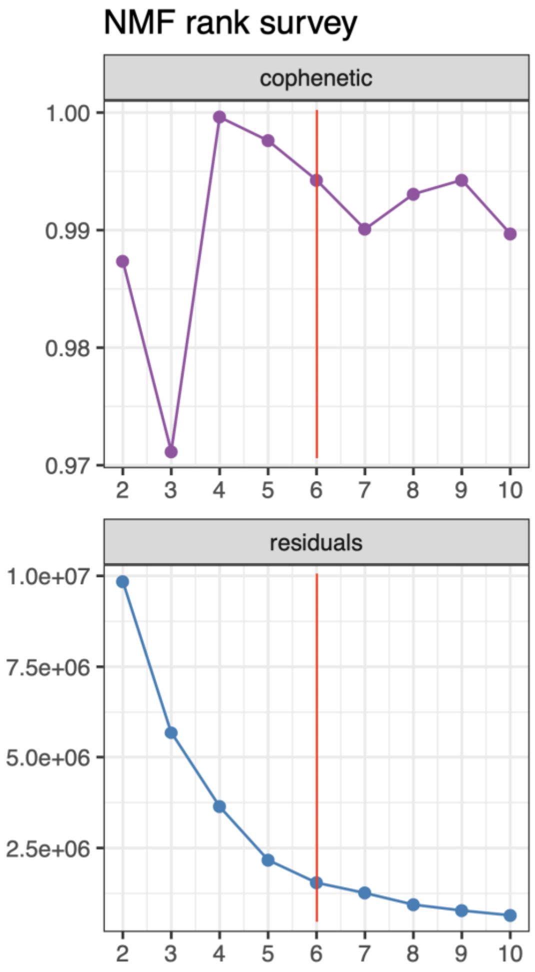


**Figure S9.** For determination of optimal dendrogram cuttoff (n=6, red line) for unsupervised hierarchical clustering, non-negative matrix factorization rank survey was performed, considering all differentially expressed transcript levels in *elav>Tau$_{R406W}$* vs. *elav* controls, based on the joint regression model, including all longitudinal data and adjusting for age. Either cophenetic correlation coefficients (top) or model residuals (bottom) are shown versus the cluster number (x-axis).

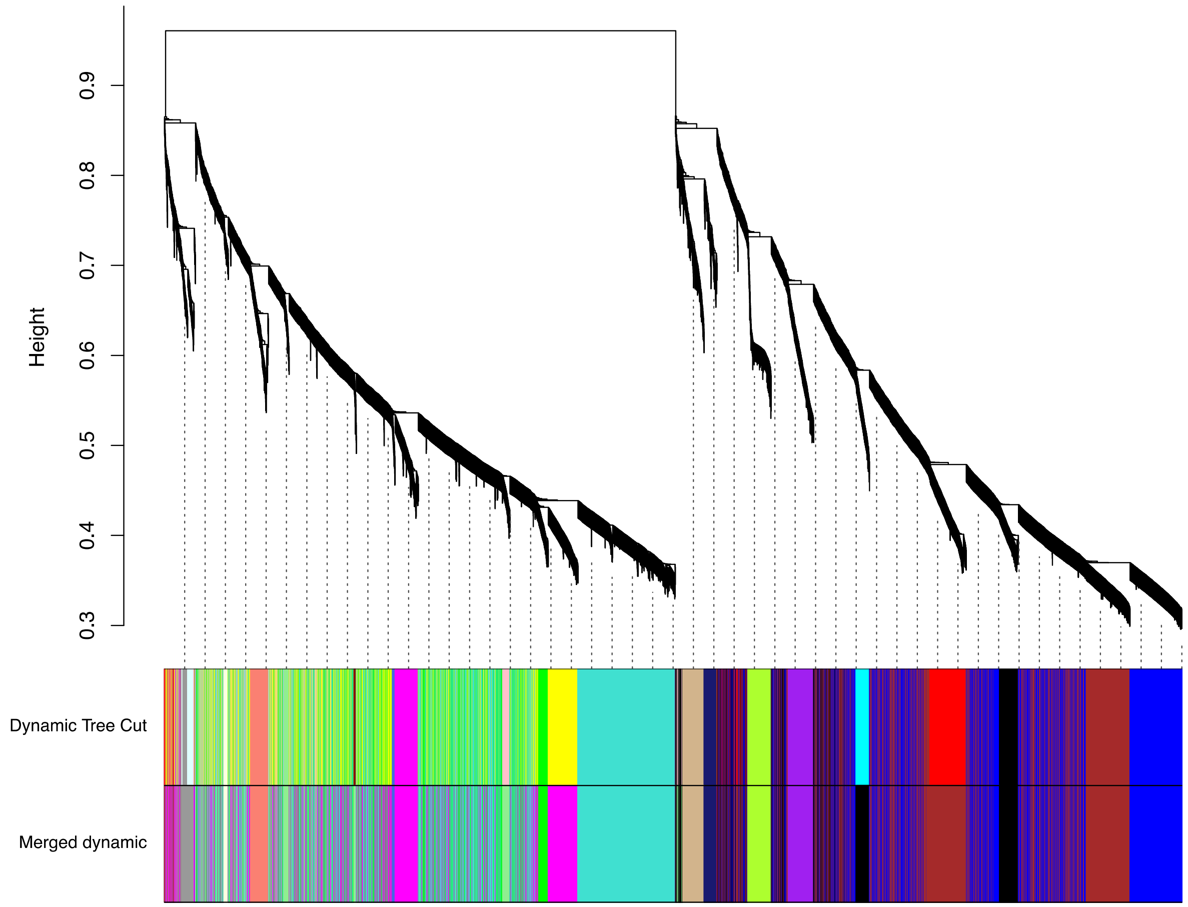


**Figure S10.** WGCNA cluster dendrogram showing tree heights/coexpression by normalized expression levels across all transcripts in *elav>Tau*$_{R406W}$ and batch-matched *elav* controls. A tree height cut was made to generate the modules shown in the color strips. The dynamic tree cut color strip denotes the modules (labeled in colors) assigned from branches in the dendrogram. The merged dynamic color strip shows new module identities after combining closely related modules (determined by module eigengene with a 0.1 tree height cutoff).
